# Characterization Eclosion Hormone Receptor function reveals differential hormonal control of ecdysis during *Drosophila* development

**DOI:** 10.1101/2025.04.03.646974

**Authors:** Valeria Silva, Robert Scott, Paulina Guajardo, Haojiang Luan, Ruben Herzog, Benjamin H. White, John Ewer

## Abstract

Neuromodulators and peptide hormones play important roles in regulating animal behavior. A well-studied example is ecdysis, which is used by insects to shed their exoskeleton at the end of each molt. Ecdysis is initiated by Ecdysis Triggering Hormone (ETH) and Eclosion Hormone (EH), which interact via positive feedback to coordinate the sequence of behavioral and physiological changes that cause exoskeleton shedding. Whereas the cell types targeted by ETH are well characterized, those targeted by EH have remained largely unknown due to limited characterization of the EH receptor (EHR). A gene encoding an EHR has been described in the oriental fruit fly, *B. dorsalis*, and in the desert locust, *Schistocerca gregaria*. However, little is known in these species about its expression pattern and its precise role at ecdysis, and no other insect EHRs are known. Here we analyze CG10738, the *Drosophila* ortholog of the *B. dorsalis* gene encoding EHR, and show that expressing it in cells confers sensitivity to EH. In addition, mutations of CG10738 specifically disrupt ecdysis, phenocopying the knockout of the EH gene. Together, these results indicate that CG10738 encodes the *Drosophila* EHR. As in *B. dorsalis*, EHR is expressed in the ETH-producing Inka cells; in addition, it is expressed in many known targets of ETH, including the neurons responsible for the secretion of other ecdysis-related peptides, such as CCAP and EH itself. Our results from targeted knockdown and rescue experiments reveal that EHR is required for ecdysis in diverse cell types and that the role of EHR in different targets differs with developmental stage. Our findings indicate extensive convergence of EH and ETH signaling and provide an exemplar of the complex mechanisms by which hormones control animal behavior.

**AUTHOR SUMMARY:** Hormones and neuromodulators are important regulators of animal behavior. In insects, one of the best studied behaviors influenced by hormones is ecdysis, which allows the animal to shed the remains of its exoskeleton at the end of each molt. Ecdysis is controlled by two key hormones: Ecdysis Triggering Hormone (ETH) and Eclosion Hormone (EH). Whereas most targets of ETH have been identified, those of EH have remained largely unknown due to the limited characterization of its receptor, EHR. Previous studies identified a gene encoding EHR in the oriental fruit fly, *Bactrocera dorsalis*, and in the desert locust, *Schistocerca gregaria,* but little is known about its expression pattern or its precise role. Here, we show that CG10738 encodes the *Drosophila* EHR. We found that EHR is expressed in ETH-producing Inka cells, in neurons that secrete other ecdysis-related peptides, in addition to other neuronal classes and non-neuronal cells. Targeted knockdown and rescue experiments revealed that EHR is essential for ecdysis in various cell types, and that its role can vary depending on the developmental stage. Our findings reveal that the role of EH at ecdysis is complex and provides insights into how hormones and neuromodulators regulate animal behavior.

## INTRODUCTION

Peptide hormones and neuromodulators are one of the most diverse groups of signaling molecules found in animals [1]. These small chains of amino acids have diverse functions both within and outside the central nervous system (CNS) in regulating a wide range of processes that that include metabolism, stress responses, circadian rhythms, locomotion, sleep, feeding, learning, memory, and social interactions [2–10]. They also play a role in regulating developmental processes such as insect metamorphosis, growth, and reproduction [11]. Through their multifaceted roles, peptide hormones and neuromodulators have profound effects on the physiology and behavior of organisms, highlighting their significance as essential modulators of neural communication [12, 13]. Determining the mechanisms by which peptides control behavior is challenging due to the complexity of their actions. Indeed, they operate at different levels, involving various signaling pathways and cellular processes. They can act rapidly or over prolonged periods, and they may exert localized or widespread actions, affecting the entire CNS, specific neurons, as well as targets outside the CNS [13–17]. The synergistic actions of multiple peptides in interconnected networks further complicates efforts to understand their complex roles in behavior and physiology.

Ecdysis provides a unique system in which to investigate the mechanism of peptide action and the complex neuromodulation of a stereotyped behavior. This essential sequence of behavioral and physiological events occurs in all arthropods, where it is used to shed the remains of the old exoskeleton at the completion of each molt, and to then inflate, harden, and pigment the exoskeleton of the next stage. This innate behavior includes three distinct and highly stereotyped phases: pre-ecdysis, ecdysis, and post-ecdysis, each characterized by specific timing and coordinated movements [18, 19]. Any disruption or alteration in these ecdysis phases can have severe consequences including death.

Ecdysis is controlled by a complex interaction of neuropeptides and peptide hormones acting in a sequential order on the CNS [18, 20, 21]. Our current understanding of the endocrine bases of ecdysis proposes that ecdysis is initiated by the release of Ecdysis Triggering Hormone (ETH) from epitracheal cells (Inka cells) and of Eclosion Hormone (EH) from the ventromedial neurons (Vm) in the brain. These two peptides trigger each other’s release, establishing an endocrine positive feedback that causes the sudden and near-complete release of both peptides [22–24] and provides an unambiguous, all-or-nothing, endocrine signal that triggers ecdysis. ETH and EH then act within the CNS, and previous work has provided information on the identity of many of the neurons that express the ETH receptor (ETHR) as well as on their timing of activation [25, 26]. Thus, ETH acts on neurons that express the neuropeptides, Crustacean Cardioactive Peptide (CCAP), Myoinhibitory Peptide (MIPs), Bursicon, FMRF-amide (FMRFa), and Leucokinin, as well as on a large number of other peptidergic and non-peptidergic targets [19, 26, 27]. Although it is known that the actions of EH are not limited to causing ETH release but that it also acts within as well as outside of the CNS to control ecdysis [28, 29], the identity and function of the EH targets are not known.

Here, we focus on the role of EH at ecdysis by identifying and characterizing the neurons and cells that express the EH receptor (EHR) in *Drosophila*. First, we confirmed that EHR is encoded by the CG10738 gene, the ortholog of a gene in *B. dorsalis* previously identified as an EHR [30]. We then used the Trojan exon technique [31] to generate an EHR-GAL4 driver and Split Gal4 hemidrivers, which we used to determine the expression pattern of the receptor and the function of EH targets as well as their pattern of activation during ecdysis. As expected, EHR is expressed in ETH-producing Inka cells. In addition it is expressed in the Vm neurons, which produce the EH neurohormone, suggesting that EH may participate together with ETH in an endocrine feedback mechanism that regulates EH release. EHR is also expressed in CCAP neurons, as well as many other neurons that express ETHR, revealing a convergence of the EH and ETH signaling systems. We found that disabling EHR function, silencing, or killing all EHR-expressing cells, was lethal at larval ecdysis, phenocopying the deficits of *Eh* null mutants [28]. By contrast, in some cases silencing EHR expression in specific subsets of cells only affected pupal or adult ecdysis, suggesting the existence of stage-specific EH targets or of targets that are essential only to ecdysis to a particular stage. Finally, we found that restoring EHR function in subgroups of EHR-expressing cells rescued normal ecdysis only partially and in a stage-dependent manner, suggesting that normal ecdysis may require EHR expression in diverse cell types, whose relevance may vary across different molts. Our findings underscore the complexity of the neuroendocrine control of ecdysis and enhance our understanding of the mechanisms by which neuropeptides function and modulate animal behaviors.

## RESULTS

### CG10738 gene function is required for EH-induced ETH release from Inka cells

Previous research in the Oriental fruit fly, *Bactrocera dorsalis*, provided strong evidence that a gene homologous to the *Drosophila* CG10738 gene encodes an EH receptor expressed in the Ecdysis Triggering Hormone (ETH)-producing epitracheal “Inka” cells [30]. To determine whether this receptor guanylyl cyclase has EHR activity, we determined whether genetic lesions in CG10738 (see Fig. S1) affected the ability of synthetic EH to trigger the release of ETH from “Inka” cells *in vitro*, which are known EH targets [22]. As shown in Fig. 1A and quantified in Fig. 1F (see also Table S3), we found that trachea from control (*w^1118^*) 1^st^ instar larvae at the dVP stage (double Vertical Plate; approximately 25 min prior ecdysis to the 2^nd^ instar; [32]) challenged with 1 nM of synthetic EH released ETH from epitracheal cells in amounts similar to the near complete depletion of ETH that is seen in control animals at the end of ecdysis (Fig. 1C). By contrast, no such release was observed (Fig. 1B) in trachea from larvae *trans*-heterozygous for a genetic deletion, *Df(3L)Exel9017* (called here *Df(3)EHR*) which includes CG10738, and either a *Minos* insertion within exon 4 of CG10738 (called here *Mi*{*EHR*}) or a Trojan GAL4 driver in which GAL4 was inserted in the intron between exons 10 and 11, which was designed to interrupt the CG10738 gene (called here EHR-GAL4)(Fig. S1). A similar result was obtained in intact mutant animals examined 2h after dVP stage (Fig. 1D and quantified in 1G; see also Table S3).

**Figure 1:**
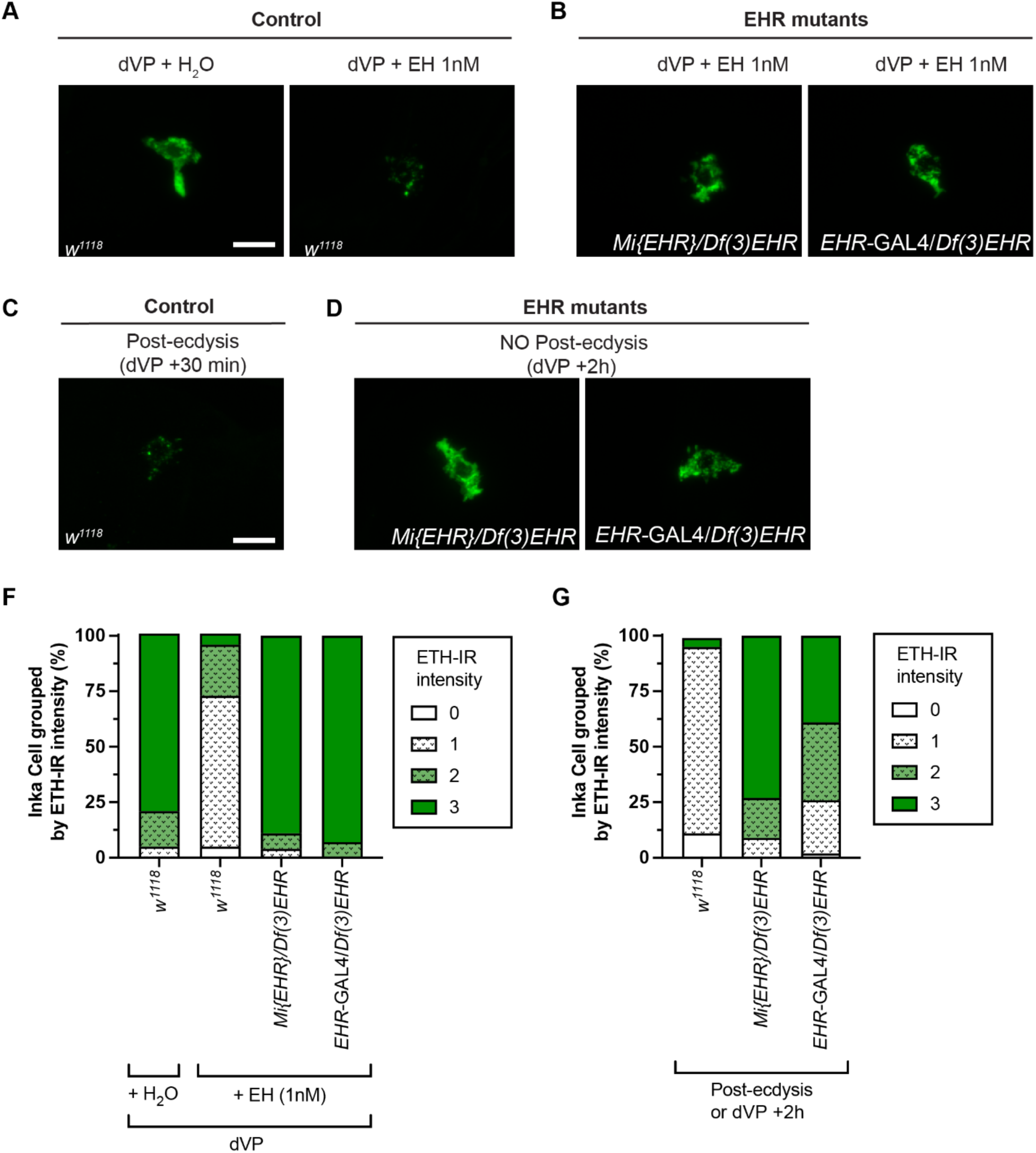
Genetic lesions in CG10738 abolish EH-induced ETH release. (A) Immunoreactivity to ETH (ETH-IR) of epitracheal Inka cells from control animals (*w^1118^*) before (left) and after (right) *in vitro* stimulation with synthetic EH. (B) ETH-IR of epitracheal cells of EHR mutant animals*, Mi*{*EHR*}/*Df(3)EHR* (left) and *EHR*-GAL4/*Df(3)EHR* (right), after *in vitro* stimulation with synthetic EH. (C) ETH-IR of epitracheal cells of wildtype (*w^1118^*) animals after ecdysis. (D) ETH-IR of epitracheal cells of EHR mutant animals, *Mi*{*EHR*}/*Df(3)EHR* (left) and *EHR*-GAL4/*Df(3)EHR* (right), 2h after dVP stage. (F, G) Quantification of ETH-IR intensity for experiments depicted in (A) and (B), and (C) and (D), respectively (N: 30-60 ETH cells, corresponding to 3-7 larvae per group). Data were analyzed by Chi-square test (see Supplementary Table S3). Scale bar 10μm.

### Mutations in EHR cause lethal phenotypes during ecdysis behavior

We next evaluated the behavior of animals *trans*-heterozygous for the *Minos* insertion in CG10738 or the GAL4 insertion in CG10738, and *Df(3L)Exel9017*. As summarized in Fig. 2, these mutant animals expressed at ecdysis phenotypes similar to those observed in *Eh* null animals [28]. Indeed, they spent at least four times longer performing ecdysis-like behaviors compared to controls (*w^1118^*); they also did not express the pre-ecdysial phase of the behavior, and the majority failed to shed their old cuticle, often dying trapped within the old exoskeleton, and expressing the “buttoned-up” phenotype (Fig. 2A, B; see also Table S3) first described for ETH mutants [32]. Overall, approximately 80% of EHR mutant larvae failed to survive the first ecdysis (Fig. 2E). Although most mutant larvae (∼70%; Fig. 2C) completely filled their trachea with air, they took approximately 25 min to do so in contrast to approximately 1 minute for control animals (Fig. 2D). The ca. 20% of animals of either EHR mutant genotype that succeeded in shedding their old cuticle eventually all died during the 2nd to 3rd larval ecdysis or during the pupal ecdysis. Importantly, both the behavioral and the tracheal defects of these mutant combinations were substantially rescued by expressing CG10738 in the EHR-expressing cells using a Trojan exon GAL4 driver inserted in gene CG10738 (Fig. 2; Table 2 and Table S3). The lethal phenotype exhibited by CG10738 mutants during ecdysis (Fig. 2) and the requirement of EHR function in ETH cells for the release of ETH upon stimulation with synthetic EH (Fig. 1), provide direct evidence that CG10738 encodes the *Drosophila* EH receptor (EHR). The similarity of the EHR mutant phenotypes to those exhibited by animals lacking a functional EH gene further suggests that this is the only EH receptor in *Drosophila*.

**Figure 2:**
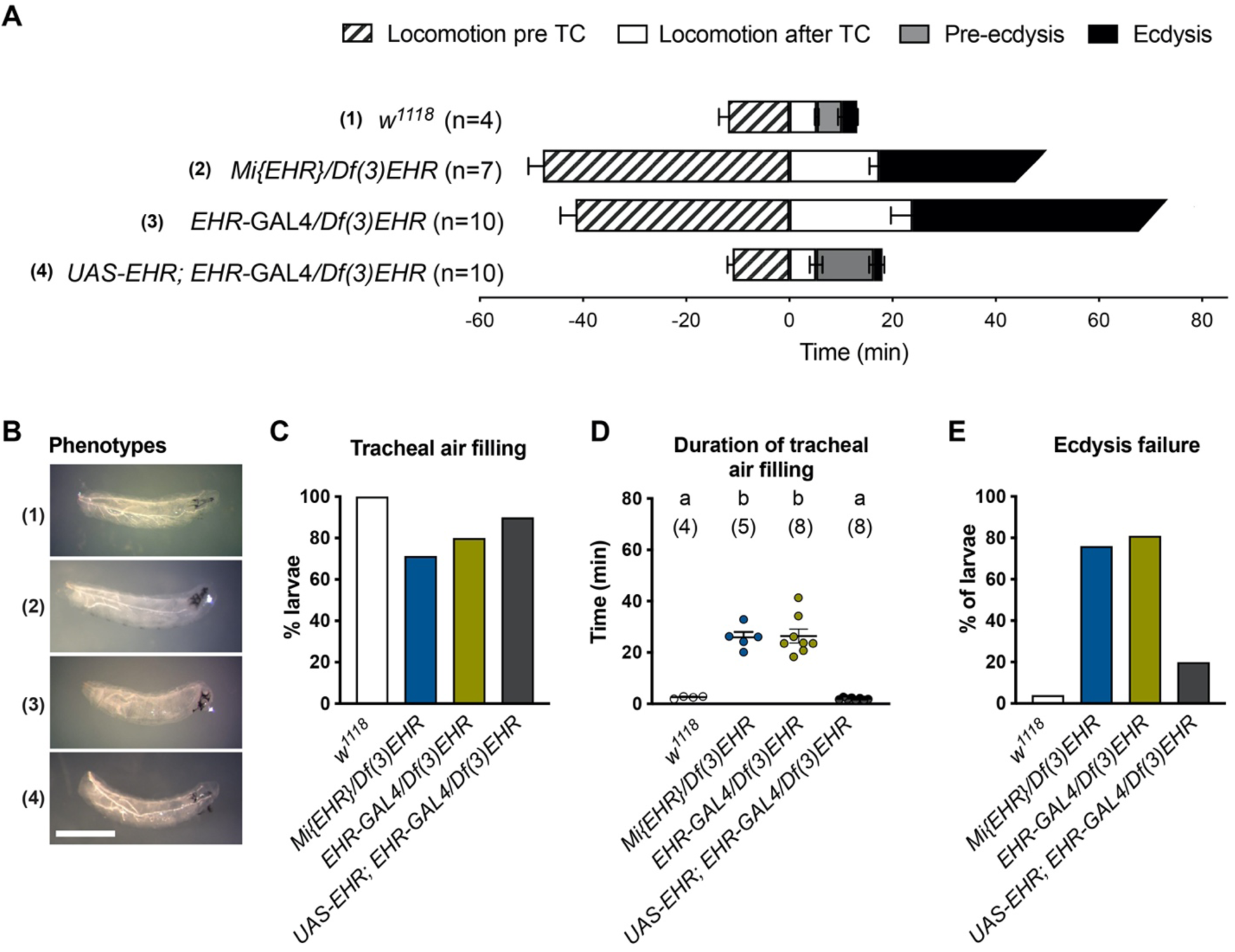
EHR mutant animals fail at the first larval ecdysis. (A) Duration of the different phases of L1 to L2 larval ecdysis. Timeline has been aligned with respect to the time of tracheal collapse (TC). (B) Representative images of animals after ecdysis (for controls) or their terminal phenotype (for EHR mutants) for (1) Control (*w^1118^*), (2) *Mi{EHR}/Df(3)EHR*, (3) *EHR-*GAL4/*Df(3)EHR*, and (4) UAS-*EHR; EHR-*GAL4/*Df(3)EHR* (genetic rescue) animals. (C) Percentage of animals that completely filled their trachea with air. (D) Duration of tracheal air filling (for animals that completed this process; N in parentheses). (E) Percentage of animals that failed larval ecdysis (N: 4-10 larvae per condition). Data were analyzed by unpaired *t*-test (A) one-way ANOVA and Tukey multiple comparison (D). Statistically significant differences are indicated by different letters in D (see Supplementary Table S3). Scale bar in (B) 500μm.

### CG10738-expressing cells are necessary for larval, pupal, and adult, ecdysis

To determine the role of cells expressing CG10738 during larval, pupal, and adult ecdysis we first examined the consequences of inactivating or ablating all EHR-expressing cells using the inwardly rectifying potassium channel, *Kir2.1* [33], or the apoptotic inductor, *reaper* [34], respectively. In both cases we observed that virtually all larvae exhibited severe ecdysis deficits, including an extended locomotory phase, an absent or atypical pre-ecdysis phase, and an ecdysial phase that lasted at least 10 times longer than normal and was ultimately unsuccessful (Fig. 3A, D; cf. Table S3), with larvae dying trapped within their old cuticle with severe deficiencies in tracheal air filling (Fig. 3B-C). We also used the TARGET system [35] to conditionally inactivate the CG10738-expressing cells 24h before pupal and adult ecdysis and found that this also caused a lethal phenotype during the corresponding ecdysis (Fig. 3E, F). Similarly, activation all the EHR-expressing cells using the temperature-sensitive cation channel, *TrpA1*, for two hours prior to pupal or adult ecdysis caused 100% lethality (Fig. S2). These results demonstrate that CG10738-expressing cells are essential for larval, pupal, and adult ecdysis. They also suggest that the timing and the sequence of activation of EH targets is critical for the proper orchestration of ecdysis.

**Figure 3:**
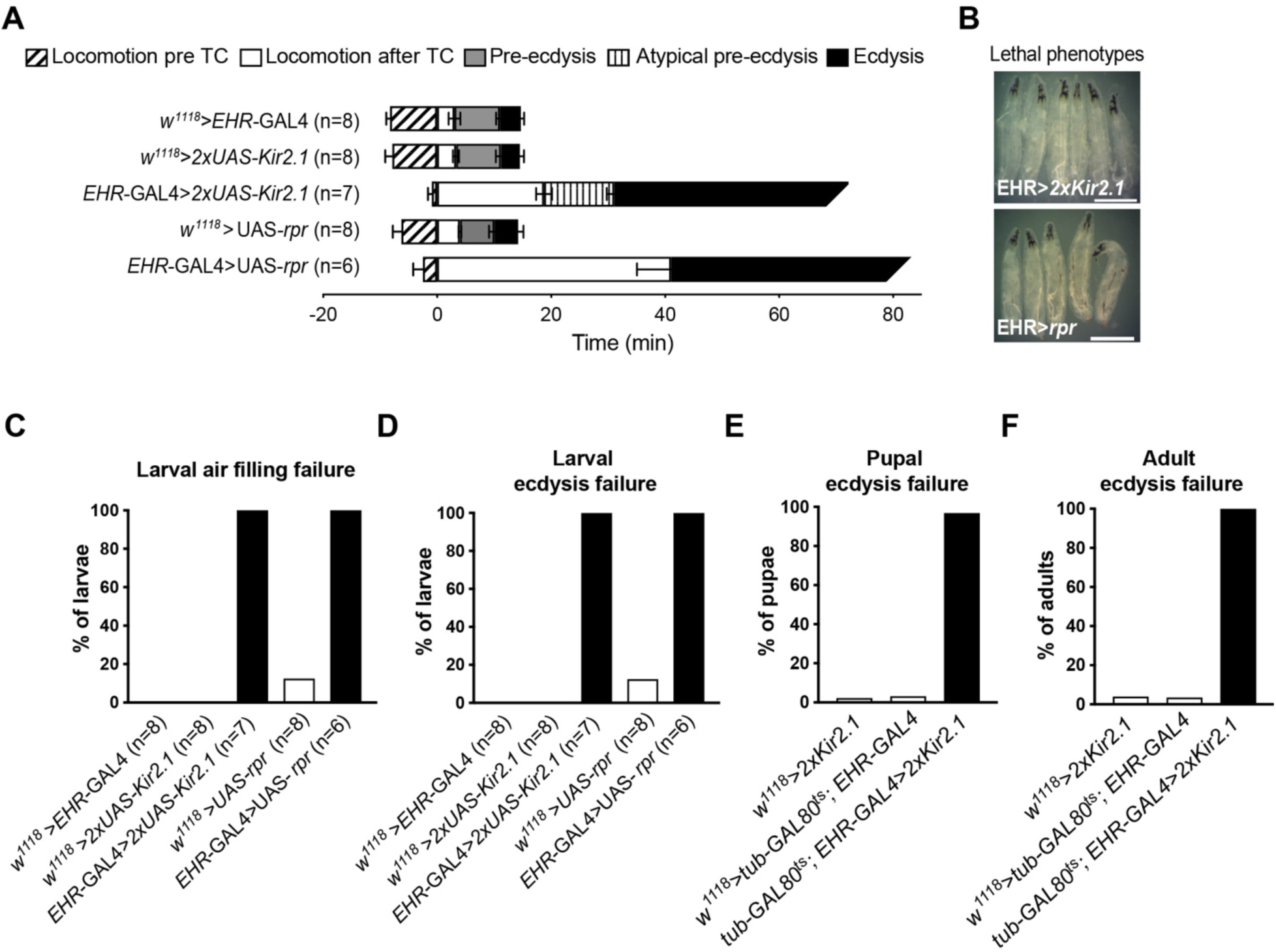
Ablating or inactivating EHR-expressing cells causes failures at larval, pupal, and adult ecdysis. (A) Duration of the different phases of L1 to L2 larval ecdysis; timeline has been aligned with respect to the time of tracheal collapse (TC). (B) Representative images of post-ecdysis or terminal phenotype of *EHR*>2x*Kir2.1* and *EHR*>*rpr* animals; (C) Percentage of animals that failed to fill their trachea with air; (D-F) percentage of animals that failed larval (D), pupal (E), and adult (F), ecdysis. (N: 150-500 per group for E-F). Data were analyzed by unpaired *t*-test (A) (see Supplementary Table S3). Scale bar 500μm

### EHR is expressed in ETH-producing Inka cells, in tracheal cells, in the CNS, as well as in other tissues, during all developmental stages

We used EHR-GAL4 to identify cellular targets of EH. Importantly, we found that EHR is expressed in ETH-producing Inka cells (Fig. 4A-C), consistent with our previous results showing that EH-induced ETH release from these cells requires EHR function (Fig. 1). EHR is also expressed in tracheal cells (Fig. 4D-E), where it plays an essential role in the rapid inflation of the new tracheal tubes that occurs at ecdysis (Figs. 2,3; [36, 37]). We also observed that EHR is expressed throughout all developmental stages in some body wall cells (Fig. S3C, E) and in the proboscis of the adult (Fig. S3E). Finally, we observed EHR expression in leg imaginal discs (Fig. 4F), which are associated with Keilin’s organ. Interestingly, cells that ring Keilin’s organ have been reported to express EH [29], which suggests a developmental signaling between this sensory structure and the developing leg.

**Figure 4:**
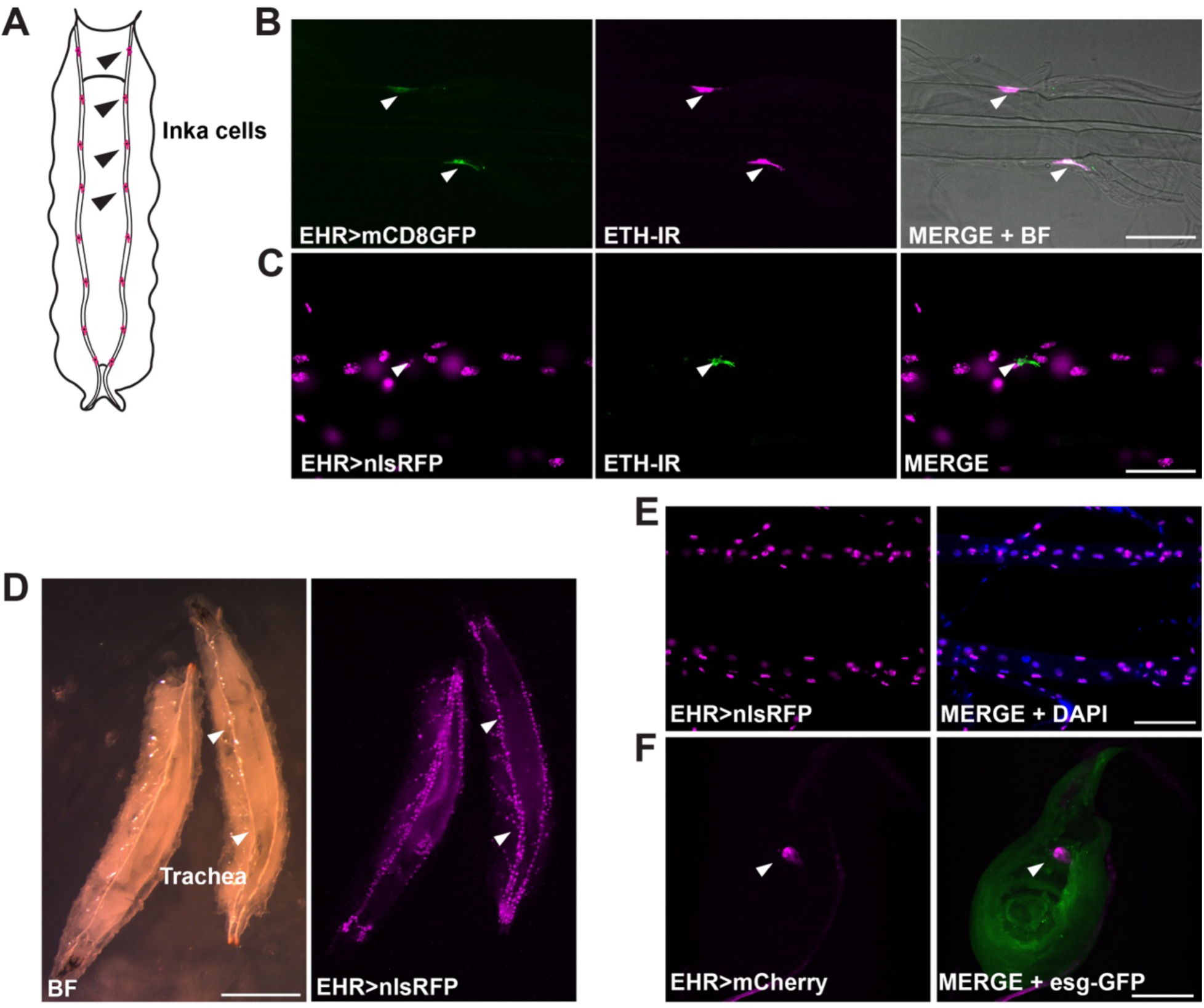
EHR is expressed in the ETH Inka cells, tracheal epithelial cells, and the leg imaginal disc of the third instar larva. (A) Schematic representation of a larva showing the Inka cells (colored in magenta; four are indicated by black arrowheads), which are located along the dorsal trunks of the trachea. (B, C) EHR expression in Inka cell (white arrowhead) visualized using the membrane-bound form of GFP (B; *EHR*>mcd8GFP) or a nuclear version of RFP (C; *EHR*>nlsRFP), together with ETH-immunoreactivity. (In B, EHR expression in tracheal cells cannot be visualized unless gain is greatly increased, probably because the cells are too large to be strongly labeled with the membrane-bound GFP used; by contrast, they are readily visible using the nuclear reporter used in (C)). (D-E) EHR expression in tracheal epithelial cells visualized using a nuclear localized RFP transgene and counterstained with DAPI (E). (F) EHR expression in leg imaginal disc, labeled using the *escargot* (*esg*)*-*GFP fusion protein. BF: bright field. Scale bars (B-C,E-F) 100μm, (D) 5mm.

We also found that EHR is broadly expressed in the larval (Fig. 5) and pharate adult (Fig. 6) CNS. In particular it is expressed in CCAP neurons at both stages (Figs. 5B, 6B), which was also expected based on work from other insects [38]. By contrast, EHR expression in EH-producing Vm neurons (Figs. 5C, 6C) was not expected, and suggests that the EH neuropeptide may participate together with ETH in controlling EH release.

**Figure 5:**
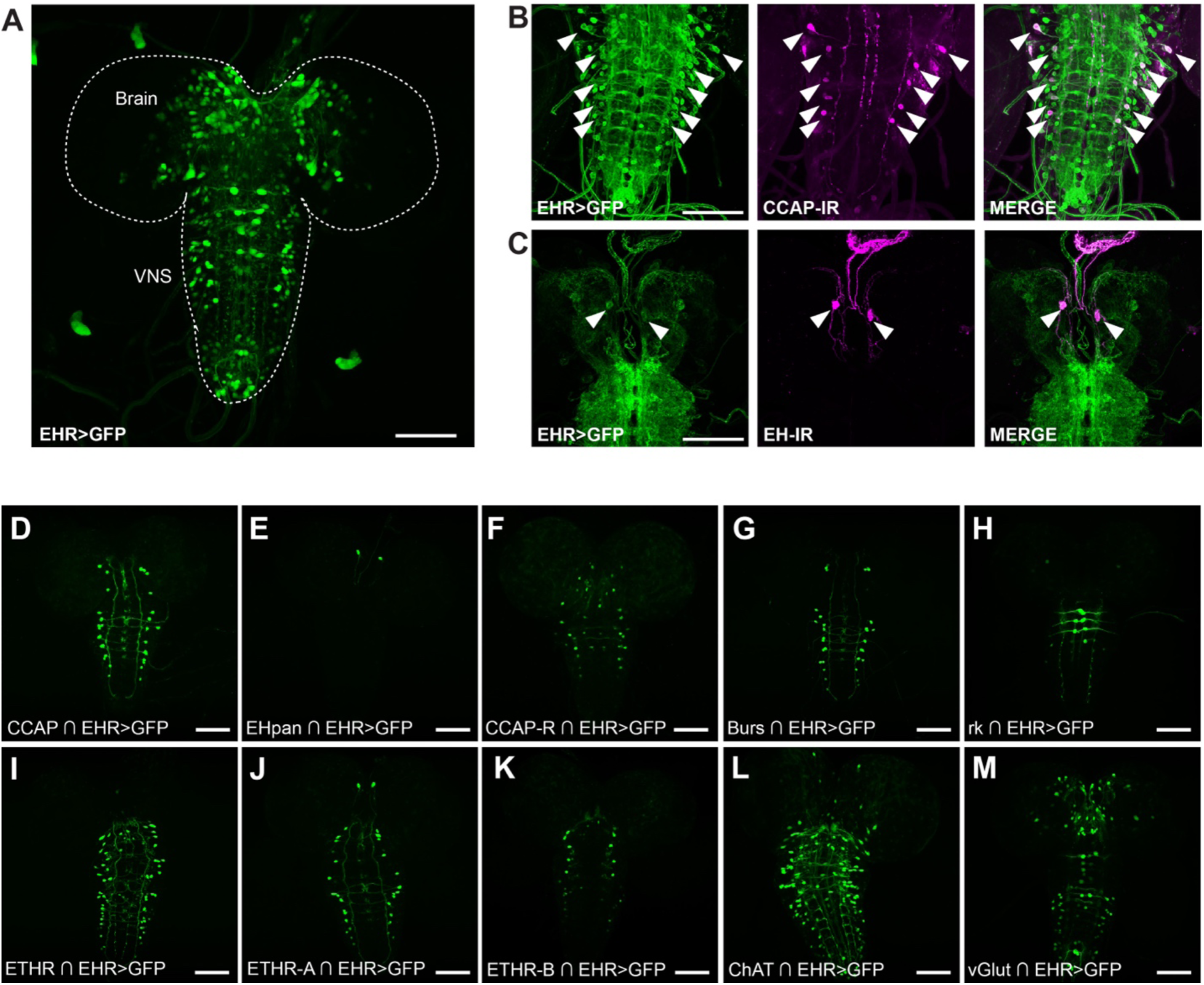
EHR expression in the larval CNS. EHR is expressed in a large number of neurons in the 3^rd^ larval instar CNS (A). In particular, EHR is expressed in CCAP neurons (B, D) and in Vm (EH) neurons (C, E). It is also expressed in neurons that express the CCAP receptor (CCAP-R) (F); in bursicon (G) and bursicon receptor (*rickets*) neurons (H); in neurons expressing the ETH receptor (ETHR) (I)(both its A (J) and B (K) isoforms); and in cholinergic (ChAT expression; L) and glutamatergic (vGlut expression; M) neurons. In (A) EHR expression was visualized using *EHR*>GFP; in (B, C), colocalization was established by co-immunolabeling with anti-CCAP (B) and anti-EH (C) antibodies; in (D-M) different subgroups of EHR-expressing neurons were visualized using appropriate “split”-GAL4 hemidrivers (indicated by intersection symbol, “∩”) together with UAS-GFP. Scale bar 100μm.

**Figure 6:**
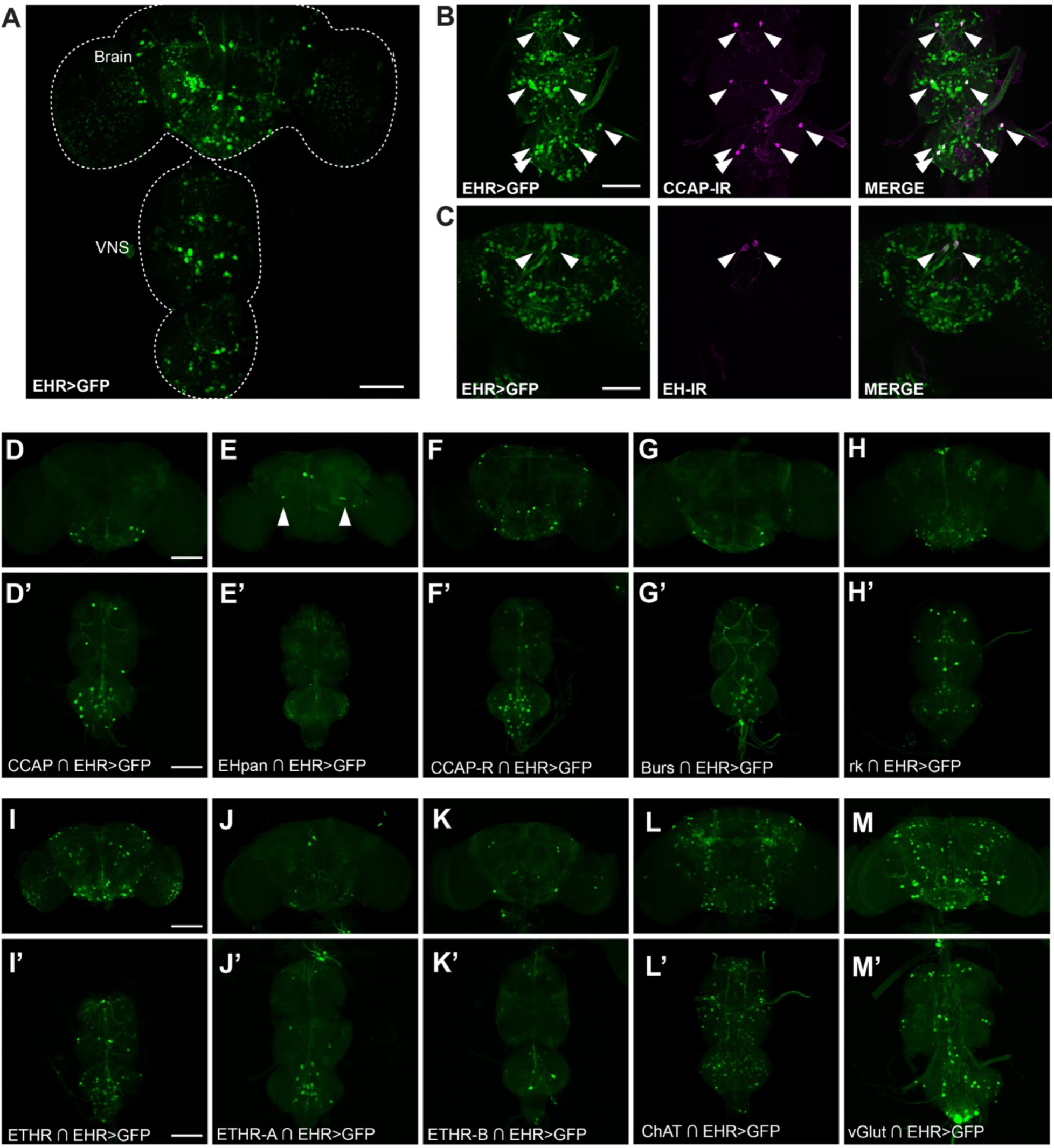
EHR expression in the pharate adult *Drosophila* CNS. EHR is expressed in a large number of neurons in the pharate adult CNS (A). In particular, EHR is expressed in CCAP neurons (B, D) and in EH neurons (Vm: C, E and Dl: E, arrowhead). It is also expressed in neurons that express the CCAP receptor (CCAP-R) (F); in bursicon (G) and bursicon receptor (*rickets*) neurons (H); in neurons expressing the ETH receptor (ETHR) (I)(both its A (J) and B (K) isoforms); and in cholinergic (ChAT expression; L) and glutamatergic (vGlut expression; M) neurons. In (A) EHR expression was visualized using *EHR*>GFP; in (B, C), colocalization was established by co-immunolabeling with anti-CCAP (B) and anti-EH (C) antibodies; in (D-M) different subgroups of EHR-expressing neurons were visualized using appropriate “split”-GAL4 hemidrivers (indicated by intersection symbol, “∩”) together with UAS-GFP. Scale bar 100μm.

We then used Split Gal4 hemidrivers to further identify neurons that expressed EHR (for these drivers, GAL4 activity is reconstituted only in cells that express both a GAL4 DNA binding domain and a transcription activation domain, thereby making them intersectional tools). Using this tool, we further confirmed EHR expression in neurons expressing CCAP (Fig. 5D, 6D,D’) and in Vm neurons (Fig. 5E; 6E, E’; arrows), both of which also express the ETH receptor [26], as well as in the Dl EH neurons of the adult (Fig. 6E; arrowheads). We also found that EHR is expressed in neurons that express the ETH receptor (ETHR, isoforms A and B) (Fig. 5I-K, 6I,I’-K,K’), thereby revealing a notable level of convergence between the signaling of EH and ETH. In addition, we found that EHR is expressed in neurons that express the CCAP receptor (CCAP-R) (Fig. 5F, 6F,F’), and the bursicon receptor, *rickets* (*rk*) (Fig. 5H, 6GH,H’), which are receptors of peptide hormones that act downstream of EH and ETH [17]. Bursicon itself is co-expressed in a subset of CCAP-expressing neurons, which, as expected, express EHR (Fig. 5G, 6G,G’). These results reveal that EH may control ecdysis through both feed forward and feedback mechanisms. Finally, the use of split GAL4 drivers showed that EH targets also include glutamatergic and cholinergic neurons (Fig. 5L-M, 6L,L’-M,M’).

### Downregulating EHR in specific subsets of EHR-expressing cells causes lethal phenotypes

In order to determine the function during ecdysis of EHR in specific subsets of EH targets, we knocked-down EHR expression in different cell types. To do so we expressed EHR RNAi under the control of different GAL4 drivers (Table 1) and evaluated the effects on larval, pupal, and adult ecdysis. We found that knockdown of EHR with the broadly-expressed *tubulin*-GAL4 driver caused significant lethality at larval stages (failure at ecdysis from 1^st^ to 2^nd^ instar and 2^nd^ to 3^rd^ instar was 47% and 95%, respectively), with the few (4) remaining 3^rd^ instar larvae failing to ecdyse to the pupal stage. Overall, few of the spatially more restricted GAL4 drivers tested caused significant ecdysial defects when used to knockdown EHR expression (Table 1). Exceptions to this were GAL4 drivers expressed in peptidergic neurons (*C929*-GAL4 which drives expression in *dimmed*-positive peptidergic neurons; [39, 40] and secretory cells (*386Y*-GAL4; [41]), the ETH cells and EH neurons, and tracheal cells (*btl*-GAL4), each of which yielded ecdysis deficits ≥ 20% at least one stage. Interestingly, the impact of the knockdown was not uniform across developmental stages, and differences occurred even between the seemingly equivalent first and second larval ecdyses. Thus, for instance, knockdown of EHR in ETH cells caused most animals (69%) to fail at the second to third instar larval ecdysis, whereas no effect was observed at the first ecdysis. Similarly, knockdown of EHR in trachea significantly affected pupal and adult ecdysis, but did not affect larval ecdysis. Whether these results reveal stage-specific differences in the control of ecdysis or are simply the result of varying knockdown efficiency is unknown. (Knockdown efficiency may vary with driver strength, which itself may vary with developmental stage. Differences in driver strength likely account for such effects as EHups-GAL4 causing more severe phenotypes than the more widely expressed EHpan-GAL4 driver driver.) Interestingly, knockdown of EHR using *n-syb*-GAL4 (neuron-specific driver) and *386Y*-GAL4 had no effect on larval ecdysis but caused 100% failure at pupal ecdysis. This result suggests that larval ecdysis may also require EH actions outside of the CNS. In this regard, it is interesting to note that somatic sources of EH have previously been implicated in the control of larval ecdysis [29]. Expression of EHR RNAi in CCAP neurons did not block eclosion but caused wing expansion defects in over half of the animals, consistent with the fact that CCAP is not essential for ecdysis in *Drosophila*, but is co-expressed with bursicon, which controls wing inflation [42, 43]. Interestingly, although there is substantial expression of EHR in ETHR positive neurons, we observed only modest defects in animals bearing knockdown of EHR in ETH targets, suggesting that EH may reinforce the response to ETH, but is not essential for ecdysial success. Finally, knockdown of EHR in CCAP-R- and *rk*- expressing cells had no overt effect on ecdysis. Thus, although the expression pattern of EHR suggests that EH may act on downstream elements of the ecdysial cascade, any such feedforward action appears not to be essential for successful ecdysis.

**Table 1:** Percentage of ecdysial failure caused by downregulation of EHR in specific subsets of cells. Drivers causing frequent failures (20% or greater) are highlighted in orange.

| GAL4 Driver | L1 to L2 | L2 to L3 | Pupal | Adult |
| --- | --- | --- | --- | --- |
| *w<sup>1118</sup>* (control) | 0% (0/101) | 0% (0/73)<sup>1</sup> | 0% (0/72) | 0% (0/93) |
| **General** |  |  |  |  |
| *tubulin* | 47% (73/156) | 95% (79/83) | 100% (4/4) | N/A<sup>2</sup> |
| *tsh* | 0% (0/24) | 0% (0/48) | 4.5% (3/67) | 0% (0/64) |
| **Peptidergic cells** |  |  |  |  |
| *c929* | 26% (7/27) | 29% (23/80) | 95% (53/56) | N/A |
| *386Y* | 1.4% (1/73) | 0% (0/108) | 100% (122/122) | N/A |
| *sNPF* | *ok*<sup>3</sup> | *ok* | *ok* | 0% (0/94) |
| *CCAP* | 0% (0/25) | 0% (0/48) | 1.7% (2/121) | 0% (0/53)<sup>4</sup> |
| *burs* | 0% (0/85) | 0% (0/112) | 0.6% (1/175) | 1.1% (2/174) |
| *EHups* | 3.3% (1/30) | 39% (19/49) | 0% (0/27) | 0% (0/27) |
| *Ehpan* | 9.4% (6/64) | 3% (2/67) | 1% (1/94) | 4.4% (4/91) |
| *ETH* | 0% (0/118) | 69% (59/85) | 100% (87/87) | N/A |
| **Neuronal classes** |  |  |  |  |
| *n-syb* | 0% (0/19) | 0% (0/37) | 100% (146/146) | N/A |
| *v-Glut* | 0% | 10% (13/130) | 0.9% (1/117) | 5.2% (6/116) |
| *ChAT* | 0% (0/41) | 0% (0/56) | 0% (0/112) | 0.9% (1/112) |
| *DDC* | 0% (0/26) | 0% (0/54) | 0% (0/46) | 0% (0/51) |
| *ple* | 0% (0/65) | 0% (0/58) | 0% (0/117) | 0% (0/151) |
| *trhn* | 0% (0/65) | 0% (0/62) | 0.6% (1/174) | 0% (0/188) |
| *GAD1* | *ok* | *ok* | *ok* | 0% (0/30) |
| *c164* | 7.1% (2/28) | 0% (0/38) | 6.4% (5/78) | 2.7% (2/73) |
| *nompC* | *ok* | *ok* | *ok* | 0% (0/74) |
| *Gr66a* |  |  | *ok* | 0% (0/146) |
| *109(2)80* |  |  | *ok* | 0% |
| *Ir20a* |  |  | *ok* | 0% |
| *Ir8a* |  |  | *ok* | 0% (0/56) |
| *Smid c161* |  |  | *ok* | 0% (0/77) |
| *Ir25a* | - | 20% (32/159) | 8.7% (11/127) | 5.2% (6/116) |
| *Ir7g* | - | 14% (42/306) | 10% (26/253) | 0.8% (2/239) |
| *Ir40a* | - | 6.9% (10/144) | 1.5% (2/134) | 3% (4/132) |
| *410-GAL4* | - | 15% (22/152) | 3.8% (5/130) | 4% (5/125) |
| **Ecdysis peptide receptors** |  |  |  |  |
| *ETHR* | 9.6% (27/280) | 13% (32/253) | 5% (11/221) | 0.4% (1/210) |
| *CCAP-R* | *ok* | *ok* | *ok* | 0% (0/23) |
| *rk* | *ok* | *ok* | *ok* | 0% (0/46) |
| **Other cells** |  |  |  |  |
| *repo* | 4.8% (2/42) | 0% (0/84) | 0% (0/94) | 7.1% (7/99) |
| *btl* | 4% (2/50) | 0% (0/52) | 54% (44/82) | 92% (35/38) |

<sup>1</sup> Fraction (and associated percentage) indicates the number of animals that failed at that ecdysis (in this case, fraction of 2<sup>nd</sup> instars that failed at ecdysis to the 3<sup>rd</sup> instar).
<sup>2</sup> No animals were obtained at this stage because an earlier ecdysis was lethal.
<sup>3</sup> The genotype of these animals could not be determined until after adult emergence, but no defects were observed at the larval or pupal stages.
<sup>4</sup> 32/53 (60%) showed abnormal wing expansion.

We were intrigued to observe EHR expression in body wall cells and in the proboscis (Fig. S3C, E) because sensory neurons play a role at *Drosophila* pupation [44], and the cibarial pump is activated at adult emergence in the moth, *Manduca sexta* [45]. Interestingly, EHR knockdown using GAL4 drivers expressed in several different subgroups of sensory neurons (Ir25a, Ir7g, and 410-Gal4) impaired ecdysial success in more than 10% of animals at least one developmental stage.

### Peptidergic neurons expressing EHR are sufficient to rescue ecdysis behavior in EHR-mutants animals

In order to identify subsets of EHR-expressing cells that may play a critical role in the control of ecdysis behavior, we investigated whether expression of EHR in particular subsets of EH targets was sufficient to rescue the ecdysial failures caused by disabling EHR function. For this, we determined whether expressing EHR using different GAL4 drivers could rescue the defects of animals mutant for EHR (mutant animals used were *trans* heterozygous for *Df(3)EHR*, and either *Mi{EHR}* or *EHR*-GAL4). These results are summarized in Table 2 and Fig. 8. As expected, expression of EHR using the EHR-GAL4 driver resulted in high levels of rescue (86-88% for larval ecdyses, and over 90% success rate in subsequent ecdyses). By contrast, most other drivers tested produced limited rescue. Overall, the most effective drivers were *C929-*GAL4 and *386Y-GAL4*, which are expressed in peptidergic or secretory cells, respectively; of these, only *C929*-GAL4 rescued adult ecdysis, albeit only modestly. In addition, the results obtained were many times stage-dependent and were greater for larval than for pupal or adult ecdyses, again suggesting that some EH targets are preferentially involved in the control of some of the fly’s ecdyses. A caveat is that just as the absolute levels of knockdown are difficult to evaluate due to the possible confounding effect of driver strength, the levels of rescue obtained using different GAL4s are also difficult to compare quantitatively. For this reason we identified drivers that produced “substantial rescue” based on whether they produced levels of rescue that were greater than those seen in EHR mutant controls.

**Table 2:**
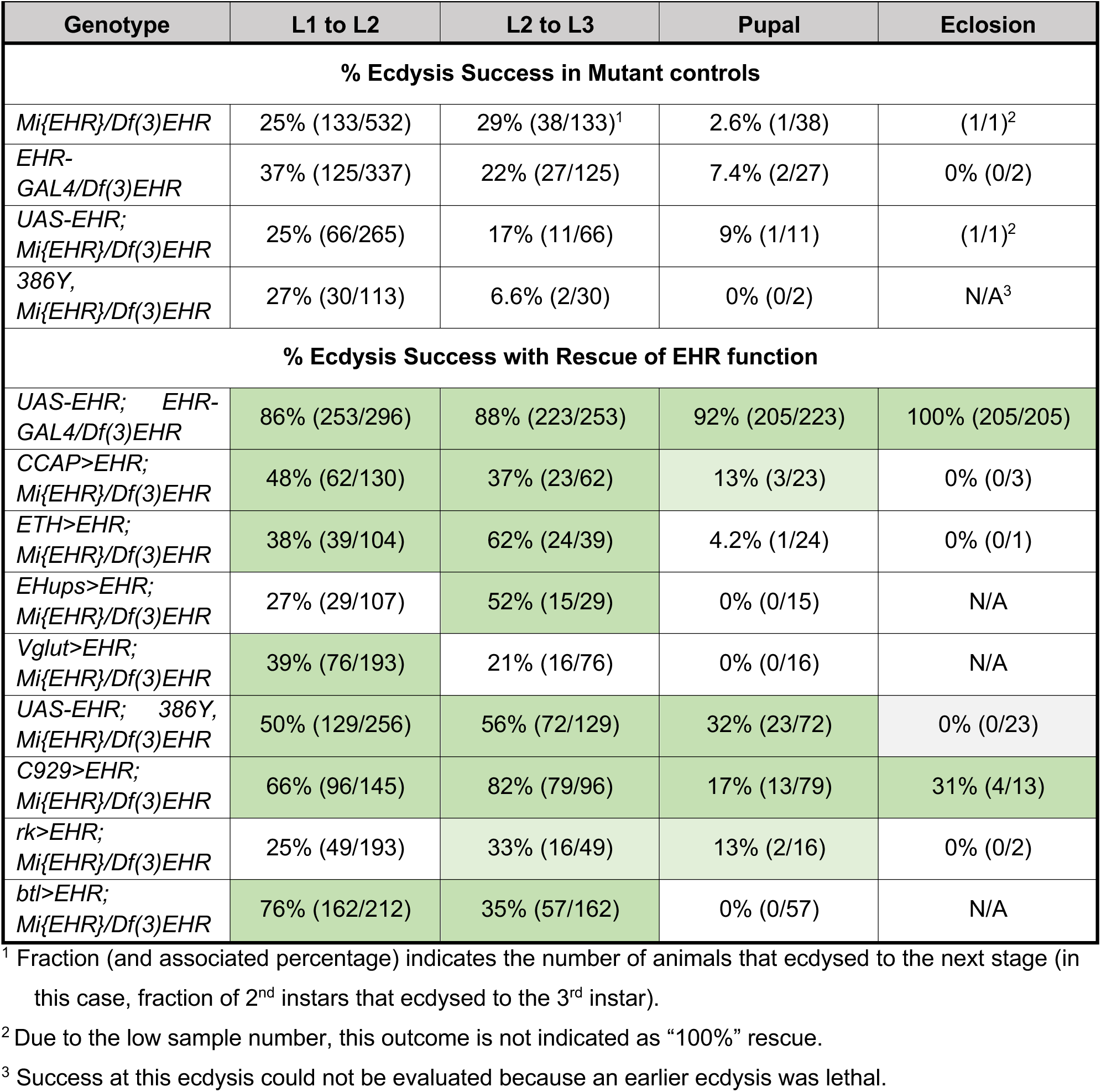
Success of genetic rescue of EHR lethal phenotype by expressing EHR in specific subsets of EHR-expressing cells. Cells marked in green indicate that a substantial rescue was observed (as compared to phenotype of the corresponding mutant control); lighter green indicates possible rescue. These data are also summarized graphically in Figure 8.

### EHR-expressing neurons show a complex pattern of activity during fictive pupal ecdysis

In order to visualize the pattern of activity induced in EH targets at ecdysis, we used synthetic ETH to stimulate the excised pupal CNS from animals expressing GCaMP6s under the control of the EHR driver (as described in [25]). Figs. 7A-C and Supplementary Video 1 shows the timecourse of activation of all individual EHR-expressing neurons after hormonal stimulation *in vitro*. We observed that most neurons were active only during the presumed ecdysial phase (min 10 to 30 of the recording), although a small group of neurons became active earlier (which might correspond to the pattern of activity of fictive pre-ecdysis) and others became active later, which might correspond to fictive post ecdysis. Thus, as has been observed previously [19, 25], different targets of ecdysial peptides respond at different times following the simultaneous stimulation with ETH, revealing that a complex pattern of neuronal activity is set in motion following the sudden release of the triggering neuropeptides, ETH and EH.

**Figure 7:**
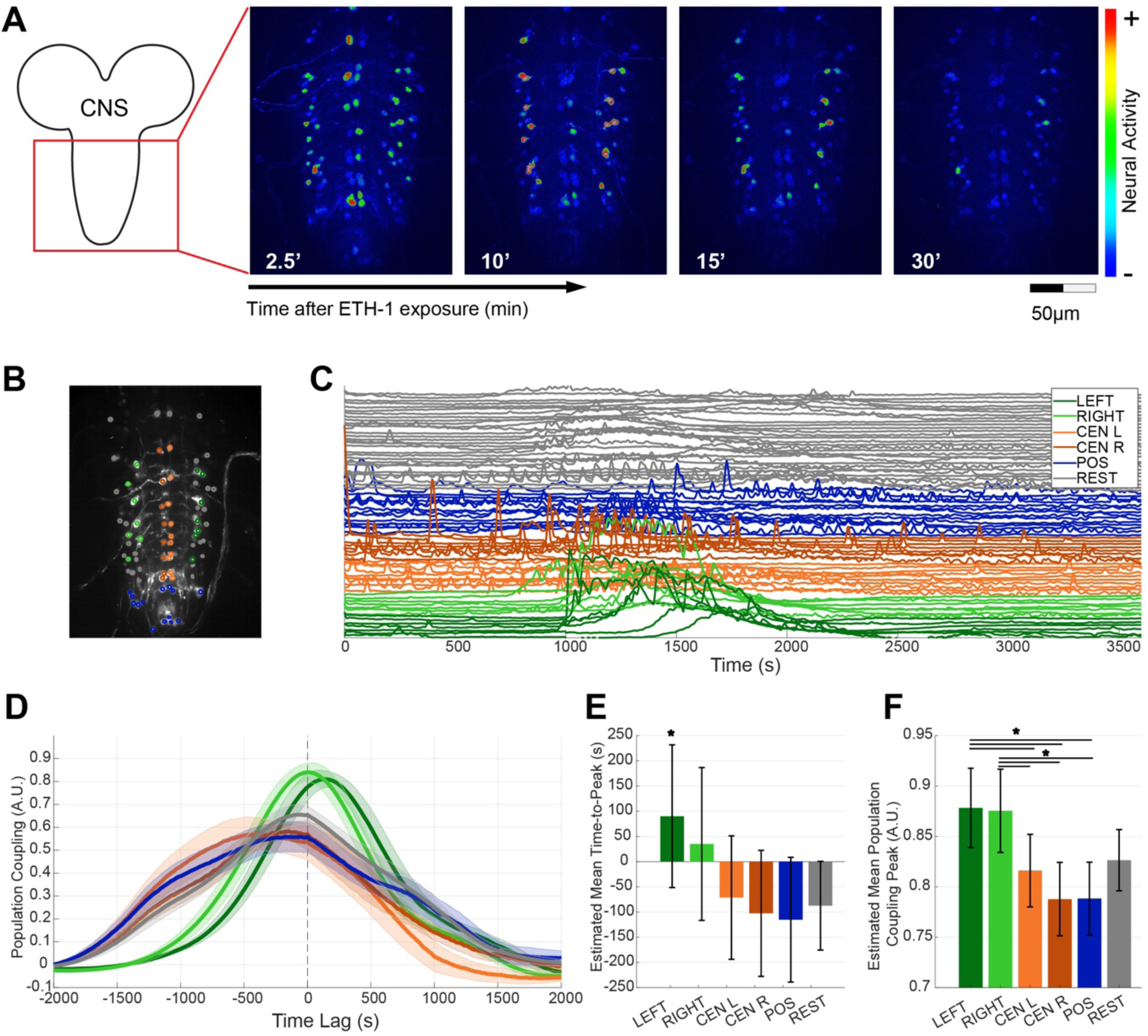
Functional population analysis of the time course of activation of EHR neurons in response to ETH stimulation *in vitro*. (A) *Left:* Reference image of a pharate pupal CNS showing the area where the CNS response was recorded. *Right*: representative frames of GCaMP signal recorded *in vitro* from CNS expressing EHR>GCaPM6s, captured at different times after ETH challenge. CNSs were imaged for 1h, with images captured at a frequency of 0.2Hz. (B-F) Functional population analysis. (B) CNS with different subpopulations of EHR neurons color coded based on their position: “Left” (dark green) and “Right” (light green), which are likely made up mostly of CCAP neurons; “Central Left” neurons (orange); “Central Right” neurons (dark orange); “Posterior” neurons (“POS”, blue); and the remaining (“REST”) neurons (gray). (C) Timeseries of GCaMP6s signal following the same color code used in B. Note the increased calcium activity around ∼1000 s (15 min) after stimulation with ETH (see also Supplementary Video 1). (D) Population coupling function for the 6 different cell types for the CNS shown in (B) and (C). Solid lines represents the average for all the cells of the same type and shaded area is the standard error. Positive lag values means that the population activates after a given cell. (E) Estimated mean time to peak of the functional population coupling function (PCF) from 5 different experiments. Asterisk denotes statistically significant differences (* p<0.05) estimated using a linear-mixed effect model. (F) Estimated mean peak of the PCF. Horizontal lines denote significant differences using Bonferroni test corrected Wilcoxon rank sum (* p<0.05) (see details in Table S3). Estimations of means and confidence intervals were derived using a linear-mixed effect model with experiments as random effects. Error bars for (E) and (F) are 95% confidence intervals.

**Figure 8:**
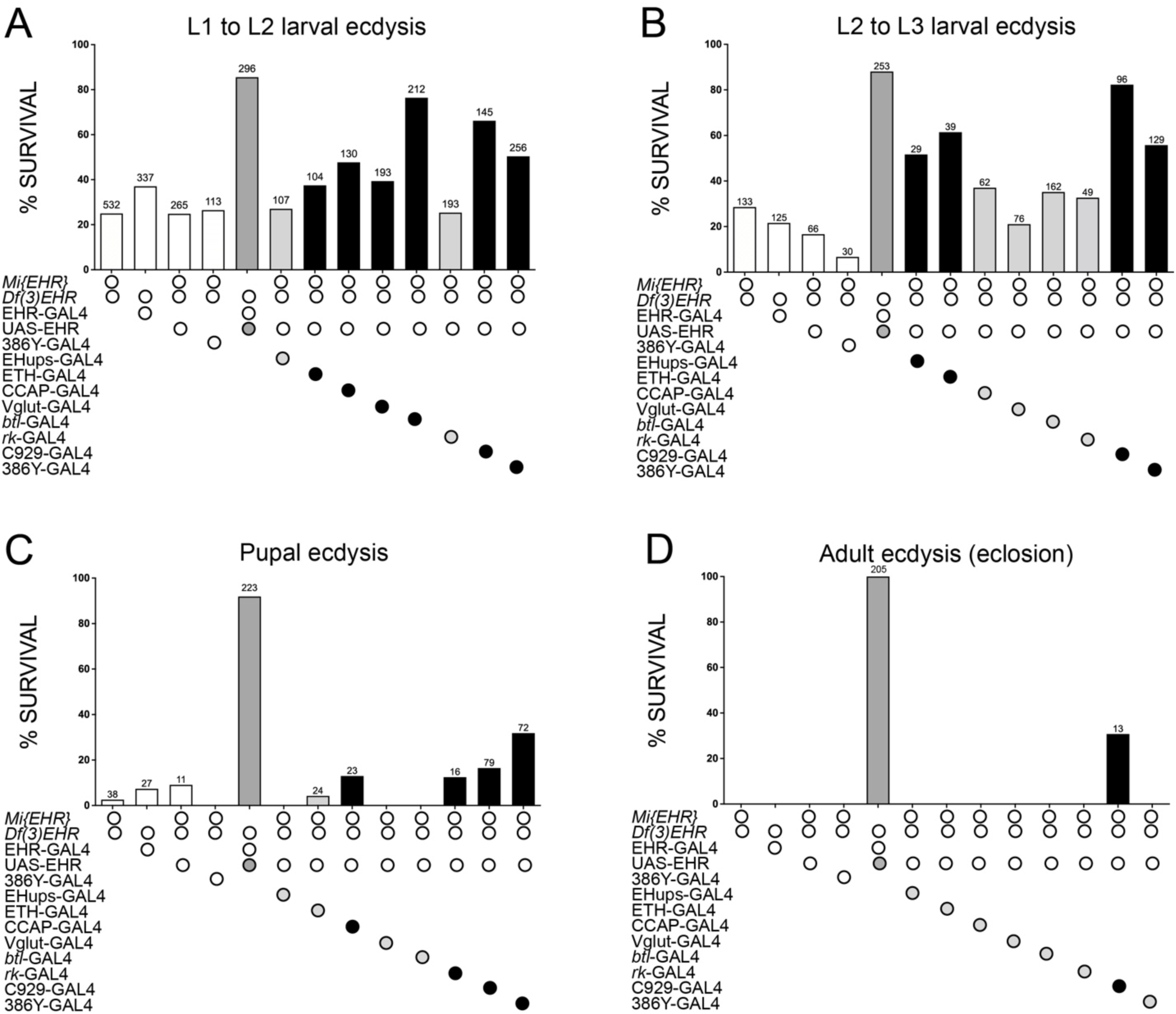
Rescue of ecdysial defects obtained by expressing EHR in subsets of EH targets. Percentage of successful ecdysis from the first to the second (A) and second to the third (B) larval ecdysis, and for pupal (C) and adult (D) ecdysis, when EHR was expressed under the control of the indicated GAL4 drivers in an EHR mutant background. Open bars summarize the phenotype of the different EHR mutants used; the dark grey bars indicate the percentage of rescue obtained expressing EHR under the control of the EHR-GAL4 driver; the light gray and black bars summarize, respectively, the results obtained for genotypes that did not (light gray bars) and did (black bars) produce substantial levels of rescue compared to animals of the corresponding mutant genetic background. Numbers above bars indicate the number of animals examined for that ecdysis. Values and percentages are shown in Table 2.

### Early Activation and Elevated Functional Coupling of CCAP Cells in the EHR Population

In order to understand the temporal dynamics within the EHR-expressing cell population, we asked whether specific neuronal populations became active earlier and displayed stronger functional coupling to the overall pattern of activity. To this end, we defined six groups of neurons based on their position (Fig. 7B; Left and Right, which are likely made up mostly of CCAP neurons; Central Left [CEN L], Central Right [CEN R], posterior neurons [POS] and the remaining neurons [‘REST’]; see Methods for further details) and computed their population coupling following Okun, *et al*. (2015) [46]. For this, we calculated for each cell group and across 5 different videos, the cross-correlation function between its activity and the average activity of the rest of the population (Fig. 7D). The temporal position (Fig. 7E) indicated whether the cell became, on average, active before or after the rest of the population and the maximum of this function served as an index of population coupling strength (Fig. 7F).

Our analysis demonstrated that CCAP cells, and in particular the “Left” subgroup, reached their peak coupling approximately 90 seconds prior to the activation of the rest of the network (p<0.05, linear-mixed effect model with experiments as random effects). The “Right” subgroup showed the same trend, but without significant effect (p=0.17). Furthermore, the amplitude of the positive peak was significantly higher for both Left and Right CCAP subpopulations compared to that of all other EHR-expressing populations (Bonferroni corrected Wilcoxon rank sums *p<0.05).

These findings position CCAP cells as early activators within the EHR population, exhibiting significantly higher functional coupling that may serve to prime subsequent population-wide activity. The marked temporal precedence and enhanced coupling of CCAP cells suggest that they play a critical role in orchestrating coordinated network dynamics.

## DISCUSSION

“Eclosion hormone” activity was described almost 50 years ago [47, 48], and EH has since emerged as one of the main actors controlling ecdysis behavior in insects. However, the breadth of its action has remained largely unknown because of the limited characterization of its receptor. Here, we demonstrate that the gene CG10738 encodes the *Drosophila* EH receptor (EHR) and show that it is expressed in all the cells and neurons previously implicated in the control of ecdysis in addition to a large number of neurons whose identity is mostly unknown. Interestingly, EH targets include the principal EH-producing neurons, many ETH targets, as well as neurons that are targets of CCAP and of bursicon, which act downstream of EH and ETH. This pattern of expression suggests that EH and ETH control ecdysis via complex neuromodulatory feedback and feedforward loops. The broad importance of EH signaling in the control of ecdysis is underscored by the phase- and stage-specific effects of manipulations of EHR function in subsets of EH targets. Thus, whereas some subsets were essential for the proper execution of all ecdyses, others were required for only pupal and adult ecdysis. Interestingly, we failed to find small subsets of EHR-expressing targets that were sufficient to rescue the ecdysial defects of most individuals of a given genotype, suggesting that many of the EH targets are important for successful ecdysis. Nevertheless, our findings underscore the relevance for successful ecdysis of EHR expression in ETH cells, in trachea, and in *dimmed*-expressing peptidergic neurons (which are defined by the C929 GAL4 driver; [39, 40]).

Although the majority of neuropeptide receptors belong to the G protein-coupled receptors (GPCRs) family [49], the EH receptor belongs to the family of guanylyl cyclase receptors (GCR). Previous research by Chang *et al.*, (2009)[30] characterized the GCR, BdmGC-1, from the Oriental fruit fly, *Bactrocera dorsalis*, and showed that it mediates robust increases of cGMP following exposure to EH, when expressed in HEK cells. They also showed that BdmGC-1 is strongly expressed in the epitracheal ETH cells and that these cells respond with an increase in cGMP-immunoreactivity to an in vitro challenge with synthetic EH. Recently, Verbakel *et al.* (2024) [50] showed that knockdown of the locust ortholog of BdmGC-1 causes failures at ecdysis. Our results here confirm that the *Drosophila* ortholog of BdmGC-1, CG10738, encodes the *Drosophila* EH receptor and greatly extend what is known about EHR distribution and function.

Although EH is known to act within the *Drosophila* CNS [28], neuronal targets of EH have not been identified directly. In many insects, cGMP-immunoreactivity (cGMP-IR) increases at ecdysis in CCAP neurons [51, 52] This is also true in some dipterans, such as mosquitoes, but has not been observed in *Drosophila*, where increases in cGMP-IR have been detected in ETH producing Inka cells [36], but not CCAP neurons [52]. Furthermore, CCAP neurons (and CCAP itself) are not required for larval ecdysis in *Drosophila* [53], suggesting that other neurons (in addition to or instead of) CCAP neurons must express an EH receptor. Here we show that EHR is widely expressed in hundreds of neurons including CCAP neurons, as well as in the trachea and epitracheal cells. Why all these targets do not show increases in cGMP at ecdysis remains unknown, but may be due to the sensitivity of the anti-cGMP antibody [52].

Interestingly we found significant colocalization of EHR and ETHR within the CNS, revealing that EH and ETH not only coordinate each other’s release, but share common targets in the control of ecdysis. In addition, EHR is expressed in the principal EH-producing Vm neurons themselves and is also co-expressed with receptors to neuropeptides that have historically been viewed as acting downstream of EH (such as bursicon and CCAP). Thus, these results indicate the presence of both feedback and feedforward loops in EH signaling. Coherent feedforward loops in which both signals coordinately activate a downstream target are often used to filter transient activation by a primary signal and delay the onset of full activation [54]. Such features might underlie, at least in part, the sequential release of neuropeptides during ecdysis and the serial activation of the different ecdysis phases. In addition, we find considerable convergence of EH and ETH signaling, including at the Vm and CCAP/bursicon neurons. This convergence suggests synergistic physiological effects of the two signaling systems on downstream neurons, perhaps to dynamically regulate excitability, though this remains to be determined. In any case, the architecture of control revealed here, with EH and ETH mutually reinforcing each other’s release and coordinating each other’s downstream actions, is reminiscent of the architecture of developmental gene regulatory networks, in which transcriptional determinants establish a regulatory state and then activate downstream inductive signals [55].

In addition to the CNS, we found that EHR was expressed in the epithelial cells of the trachea, including cells from the primary tubular branch, secondary branches, and terminal branches (Fig. 4C, D), and that knockdown of EHR in trachea caused failures at ecdysis. These findings are consistent with previous reports that showed that ablating the Vm neurons [37] or disabling the *Eh* gene [28] caused defects in the vital process of tracheal air-filling that occurs during ecdysis. These defects cannot be rescued by injections of ETH [36] indicating that they are due to the action of EH and are not due to the absence of ETH release associated with these genotypes.

Although the identity of all EH targets largely remains to be determined, the relatively penetrant effects of EHR suppression in sets of peptidergic neurons, as opposed to neurons that use fast neurotransmitters (see Table 1), suggests that EH signaling is most important in modulating cells in the ecdysis network above the level of the synaptic networks that directly generate behavior. This is consistent with a model in which modulatory interactions regulate behavioral state, which is then translated into the execution of state-specific motor programs. The fact that both ecdysis motor programs and the function of subsets of EHR-expressing cells vary with developmental stage suggests that either the identity or the importance of the downstream factors recruited by EH changes over development. This plasticity in ecdysis peptide usage is also seen phylogenetically, with the complement of essential ecdysial peptides varying across insect species. For example, whereas CCAP and orkokinin are dispensable for ecdysis in *Drosophila* [53, 56], both are essential in the kissing bug, *Rhodnius prolixis* [57, 58]. The stage-dependent differences in function of both ecdysial peptides and the cell types that express them in *Drosophila* may, in fact, simply reflect necessary differences in the control and generation of the ecdysis motor programs that evolved to accommodate metamorphosis and its attendant changes in body plan [59]. Overall, our results suggest an intricate interplay between multiple cell types and between multiple neuromodulatory peptides in regulating ecdysis. Understanding of how this complex behavior is controlled in *Drosophila* and, potentially, in other species, should contribute to our understanding of how neuromodulation mediated by neuropeptides regulates animal behaviors.

## MATERIALS AND METHODS

### Fly strains

Fly stocks were maintained at room temperature (22-25°C) on standard cornmeal agar media under a 12 h light/12h dark regime. Stocks used and their source are listed in Table S1.

### EHR-GAL4 generation

The EHR-GAL4 line was made by inserting a Trojan Gal4 Expression Module (TGEM) [31] into the intron separating exons 10 and 11 of the CG10738 gene (Fig. S1). Insertion was targeted to the following site using CRISPR/Cas9: AAATGTGGGCTGTACTTATT<u>AGG</u> (PAM site underlined). To make the TGEM construct, homologous arms of 1 kb flanking the Cas9 cleavage site were amplified by PCR using the following primer pairs synthesized by Integrated DNA Technologies, Inc. (Coralville, Iowa, USA): EHR-3NotI: AGTCAG GCGGCCGC AAGTACAGCCCACATTTTGC, EHR-3AgeI: AGTCAG ACCGGT TGATGGAAGTCCTGGATTCG, EHR-5SpeI: AGTCAG ACTAGT AATAGGCATCCCCTCGTTGT, EHR-5AscI: AGTCAG GGCGCGCC ATTAGGTTATTTTAAAGTGCATGCAGAAGG, The PCR products were cloned into the pT-GEM(0) vector. The resulting plasmid (250-pGS3-AgeI-NotI-arm-SA-T2AGal4-hsp70-3xP3-RFP-SV40-AscI-EHRarm) was co-injected with a pBS-U6-sgRNA-ETH plasmid encoding the guide RNA into embryos of flies expressing germline Cas9 as described previously [31]. Transformants were identified by their expression of the 3xP3-RFP marker. The EHR Split-GAL4 hemidriver lines were generated from the EHR-Gal4 strain by ΦC31-mediated cassette exchange as described in Diao *et al.,* (2015) [31].

### UAS-EHR generation

GC10738 cDNA was amplified from *Drosophila* larval CNSs using standard methods. The entire coding sequence was re-amplified using primers that included XhoI and KpnI sites at the 5’ and 3’ ends, respectively. Correct sequence was confirmed and the XhoI-KpnI fragment (containing the entire CG10738 coding sequence) was cloned into pUAST transformation vector by Genewiz (New Jersey, USA). Transformants were generated by BestGene (California, USA).

### EH synthesis

Synthetic EH was produced by GenScript (Hong Kong) using the pFastBac baculovirus expression system. Several constructs were tested with the goal of producing a secreted protein with high yield. Best results were obtained with a design that included the putative native signal sequence (vs. the signal sequence of the major envelope glycoprotein, gp67) and a His-tag at the amino-terminal of the protein. The sequence of the expressed protein was (putative mature EH sequence underlined): MNCKPLILCTFVAVAMCLVHFGNAHHHHHHENLYFQG<u>LPAISHYTHKRFDSMGGIDFV QVCLNNCVQCKTMLGDYFQGQTCALSCLKFK GKAIPDCEDIASIAPFLNALE</u> Analysis indicated that the protein was not secreted and it was therefore purified from the cell lysate. It was then aliquoted in 50 mM Tris-HCl, 500 mM NaCl, 5% Glycerol, pH 8.0 at a concentration of 0.25 mg/ml and stored at −20°C.

### Stimulation with synthetic EH

Larvae were reared at 25°C on Petri dishes with fly medium. First instar larvae approaching ecdysis to the second instar were identified by their double mouth hooks (dMH), placed on Petri dishes coated with apple juice agar, and monitored until they first pigmented the plates of the second instar (double vertical plate stage, dVP, approximately 25 min prior ecdysis to the 2^nd^ instar; [32]. Their tracheae were then dissected in PBS and incubated in Schneider’s insect medium (Sigma). All tracheae of the same genotype dissected within 20 min were pooled; then were challenged either with synthetic EH (1nM final concentration) or the same volume of water. After 1h incubation at room temperature the tracheae were fixed and processed for ETH immunoreactivity [32], as described below. Tracheae from wild type animals prior to ecdysis as well as from larvae post-ecdysis were included as controls. In the case of animals mutant for EHR (which do not ecdyse) tracheae were dissected 2h after the dVP stage.

All epitracheal cells were imaged using the same parameters using an Olympus DSU spinning disk microscope (40x oil objective Olympus). The intensity of ETH immunoreactivity (ETH-IR) of each epitracheal cell was scored blind and assigned a score from 0 (no detectable ETH-IR) to 3 (maximal ETH-IR). Thirty to 60 cells were measured per conditions, representing the tracheae of 3-7 animals.

### Immunohistochemistry

Preparations were dissected on Sylgard-coated plates under cold PBS and fixed in 4% paraformaldehyde for 1h at room temperature or overnight at 4°C, as previously described in Clark, *et al.* (2004) [36]. Tissues were then rinsed several times in PBS with 0.3% Triton X-100 (Sigma) (PBSTX) and incubated in primary antibodies overnight at 4°C with constant agitation. Then they were washed in PBSTX and incubated in fluorescent secondary antibodies (Jackson ImmunoResearch, West Grove, PA) and mounted onto poly-L-lysine (Sigma) coated coverslips. Primary antibodies used include: rabbit anti-CCAP ([52]; used at 1:500), rabbit anti-ETH ([32] used at 1:2,000) and rabbit anti-EH ([28]; used at 1:200). For GAL4 lines that produced low GFP signal when driving UAS-GFP, tissues were processed using anti-GFP (Invitrogen; used at 1:1,000). Preparations were viewed and imaged under a conventional fluorescent microscope or under an Olympus DSU spinning disk microscope and analyzed using ImageJ [60].

### Survival across the different ecdyses

To quantify the number of animals that successfully completed larval ecdysis, we placed first instar larvae of the relevant genotype on Petri dishes with regular food and counted the number of larvae that ecdysed to the second instar. These larvae were transferred to Petri dishes with new regular food, and we then counted the number of animals that ecdysed to the third larval instar. We then transferred these larvae to vials with regular food, where we followed the success of pupal and adult ecdysis. For some genotypes we preferentially focused on specific ecdyses.

### Quantification of larval ecdysis behavior

Larvae were reared at 25°C on Petri dishes with fly medium. First instar larvae at the dMH stage were placed on Petri dishes coated with apple juice agar with yeast. At dVP larvae were transferred individually to a new slightly wet agar Petri dish and video-recorded under a Leica dissecting microscope (Nussloch, Germany) until they completed ecdysis or for up to 2h after the time of ecdysis of control animals. Behavioral analysis scored for the presence or absence of three different phases: “locomotion,” consisting of normal locomotor activity; “pre-ecdysis,” consisting of anterior-posterior contractions (AP) and squeezing waves (SW); and “ecdysis” which started with “biting” behavior and ended with the final backward thrust, regardless of whether the old cuticle was eventually shed. “Atypical pre-ecdysis” was defined by partial, weak, or missing AP contractions or SW (as described previously in Scott, *et al*. (2020) [29]. Quantification of tracheal dynamics included: (1) the time taken to completely fill the trachea with air; for those animals that successfully completed that process and, (2) the percentage of animals that completed the air filling process (if ≥ 95% of the trachea filled with air it was considered filled) or failed.

### Quantification of pupal ecdysis and adult eclosion

Flies were reared at 25°C on standard cornmeal agar media under a 12 h light/12h dark regime. Pupal and adult ecdysis survival and failure were evaluated as described in Lahr, *et al*. (2012) [53]. For experiments using the TARGET system (temperature sensitive GAL80^ts^; [35]) or the temperature-sensitive cation channel, *TrpA1* [61], flies were reared at 18°C and transferred to 30°C before pupal ecdysis or adult eclosion (24h hours before ecdysis for GAL80^ts^, and 2h for *TrpA1* flies), and ecdysis success or failure was evaluated.

### RNAi screening

Six different UAS-RNAi lines for CG10738 (see Table S2) were crossed with the *tubulin*-GAL4 line and the severity of the ecdysial defects determined by counting the number of larvae or pupae showing failures at ecdysis and the number of adults that emerged (see Table S2). This screen revealed that line CG10738 RNAi-R1 produced the most severe phenotype and is the one used here; it was always used in combination with UAS-*dicer2* (UAS-*dcr2*) to boost the effectiveness of RNAi knockdown.

### Real-time GCaMP Imaging

In order to record calcium dynamics in *in vitro* CNS preparations, CNSs were dissected from animals around 2 hours prior to pupal ecdysis under cold 1X PBS and immediately mounted on a poly-L-lysine coated glass, placed in an Attofluor chamber (Thermo-Fisher Scientific), and covered with 2 mL of Schneider’s insect medium (Sigma). GCaMP6s [62] signal was recorded using an Olympus DSU spinning disk microscope with a 20X water objective and a Hamamatsu high sensitivity camera. Preparations were first imaged for 5 min at 0.2 Hz, and preparations showing spontaneous activity were discarded. Synthetic ETH (600 nM final concentration) or H_2_O (control) was added and 5 Z-stacks across the whole VNC were recorded for one hour at 0.2 Hz, as previously described in Mena, *et al*. (2016) [25].

### Video analysis

Real-time calcium imaging videos were processed using FIJI software (Schindelin et al., 2012) to create hyperstacks and a maximum intensity projection (template). We then used the FluoroSNNAP MATLAB program to select all ROIs that corresponded to the center of each neuronal cell body and followed each of their activity during the 60 min recording period to create at the time series of fluorescence intensity (Patel et al., 2015). Calcium traces were normalized by using the previous 50 frames corresponding to 250 s as baseline, obtaining the dF/F traces (as described in [63] and [64]).

### Manual selection of cell types

We analyzed 5 different videos (n= 65.4 ± 10.78 cells; mean ± standard deviation), where we defined six EHR cell populations: posterior cells (POS, n= 10.2 ± 2.2), Left neurons (Left, n= 7.8 ± 0.40), Right neurons (Right, n= 6.8 ± 1.3), central cells Left (CEN L, n= 10.4 ± 2.4), central cells Right (CEN R, n= 10.2 ± 2.2) and the rest (“REST”, n= 20.20 ± 13). These groups were defined by identity and their position within the VNC considering a midline that symmetrically separate left and right side of the CNS.

### Population coupling function

The functional population coupling function (PCF) of each cell was computed following Okun, *et al*. (2015) [46]. We computed the lagged-cross correlation function between each time series of one cell versus that of a ‘population’ time series, which was obtained by averaging the activity of all the remaining cells of the population. To then characterize the level and temporality of the coupling of each cell, the maximum of the function and its location in terms of temporal lags were extracted. Using this measure, a positive lag indicates that the population follows a given neuron, whereas a negative lag indicates that the neuron follows the population activity.

A linear-mixed effect model with experiments as random effects was used to evaluate if the lags associated with the maximum of the PCF were significantly different from zero. One model was used for each cell type. Means were estimated from these models, using the “REST” group as baseline. A Wilcoxon rank sums test was used to perform pairwise comparisons between the maximum value of the PCF, and the p-values were corrected by the Bonferroni method (15 comparisons).

### Statistics

Statistical analyses were performed using GraphPad Prism (GraphPad Software, La Jolla, CA, USA) and visualized using Adobe Illustrator. If data were normally distributed and had equal variance, they were analyzed by one-way ANOVA followed by the Tukey test for *post hoc* multiple comparison analyses. Categorical data were analyzed by Chi-square or Fisher’s exact test. The exact values for each comparison are shown in Supplementary Table S3.

## Supporting information

Silva et al, Supplementary materials

Silva et al, Supplementary Video 1

## ACKNOWLEDGMENTS

We thank Eileen Krüger for amplifying CG10738 cDNA. We thank Vivian Budnik, Susan McNabb, Paul Taghert, Rob Jackson, Christian Wegener, and the Bloomington, Kyoto and Vienna, *Drosophila* stock centers for fly stocks, and Michael Adams for anti-ETH antisera. This work was supported by CONICYT Graduate Fellowship 21191720 (to V.S.); FONDECYT (National Fund for Scientific and Technological Development, Chile) grant 1221270 (to J.E.) and the Intramural Research Program of the National Institute of Mental Health, USA (ZIAMH002800, to B.H.W.).

