## Supplementary material for "Characterization Eclosion Hormone Receptor function reveals differential hormonal control of ecdysis during *Drosophila* development": Silva et al, Supplementary materials

### SUPPLEMENTARY FIGURES

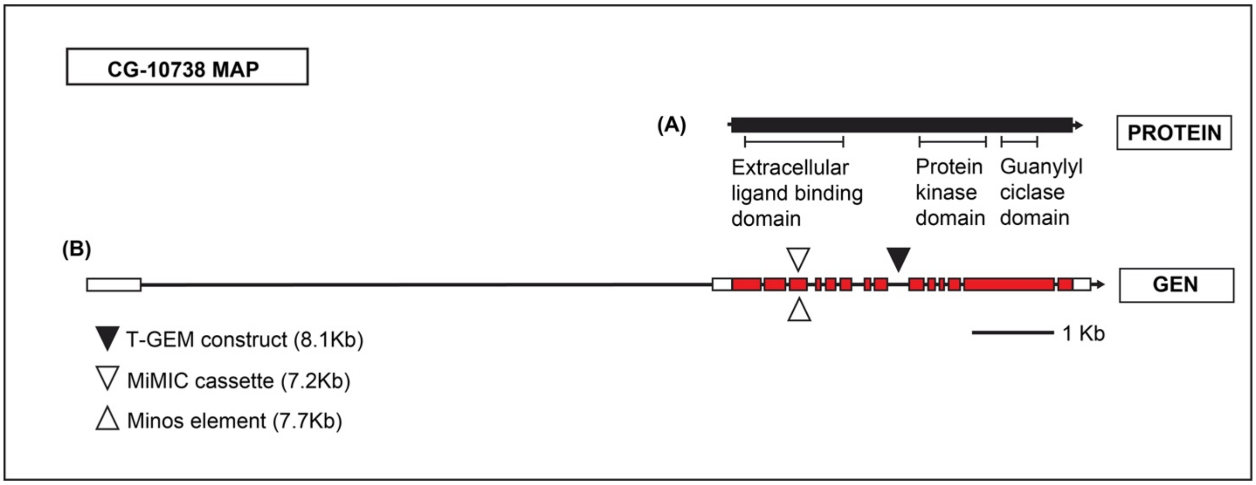


**Supplementary Figure S1: EHR protein and gene map.**

Schematic representation of the CG10738 protein (A) and gene (B). (A) Protein symbolized as a black box includes three domains predicted by InterPro. (B) Gene map shows the location of insertions used here, which include the EHR-GAL4 (T-GEM construct), and a MiMIC and a Minos insert. Non-coding regions are indicated as white boxes, coding regions as red boxes, and introns as black lines. Scale bar: 1 Kb.

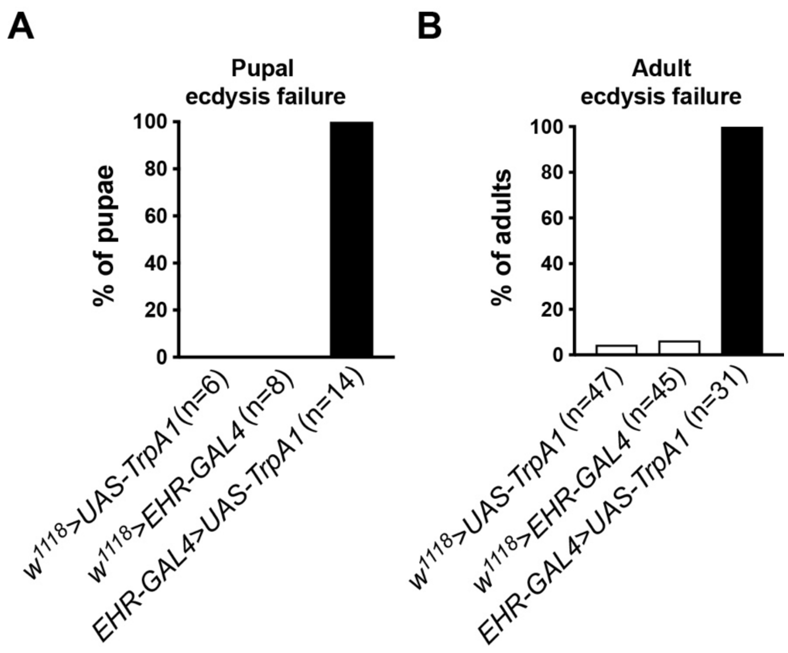


**Supplementary Figure S2: Activation of all EHR-expressing cells using TrpA1 caused failures at pupal ecdysis and at adult eclosion.**

Percentage of animals that failed to complete pupal ecdysis (A), and adult eclosion (B), after activation of EHR-expressing cells using UAS-*TrpA1* for 2h before ecdysis.

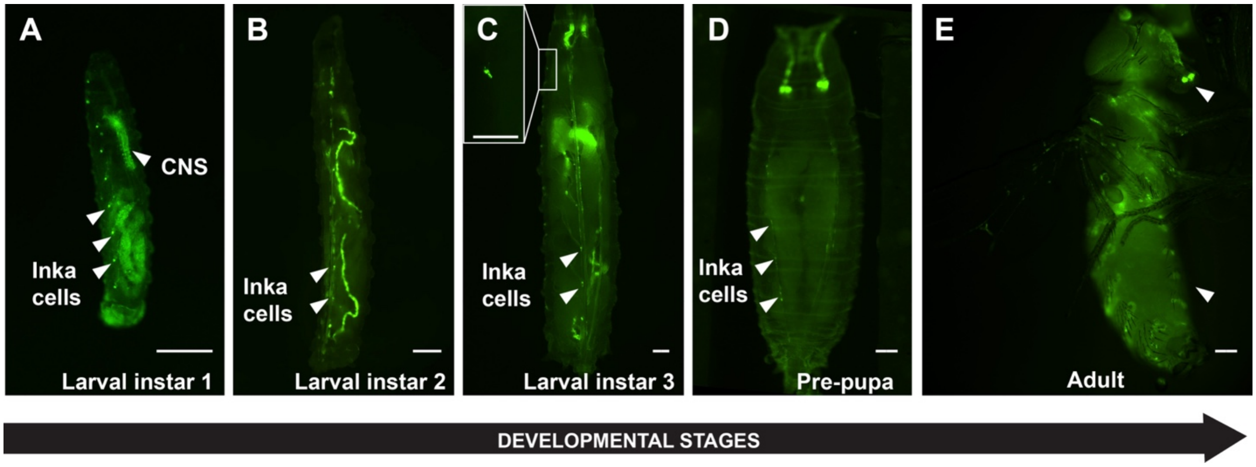


**Supplementary Figure S3: EHR is broadly expressed in somatic tissues and across developmental stages.**

Whole body pattern of EHR expression visualized with GFP (EHR>GFP) at the first (A), second (B), and third (C) larval instar; at the P3 pre-pupal stage (D); and at the pharate adult stage (E). Insert in (C) shows EHR expression in some cells of the body wall; white arrowheads in (E) point the EHR expression in the proboscis (top) and in dorsal bands of cells of the body wall (bottom). Scale bar 200 μm for panels and (A-E), and 100 μm for insert in (C).

**Supplementary Video**

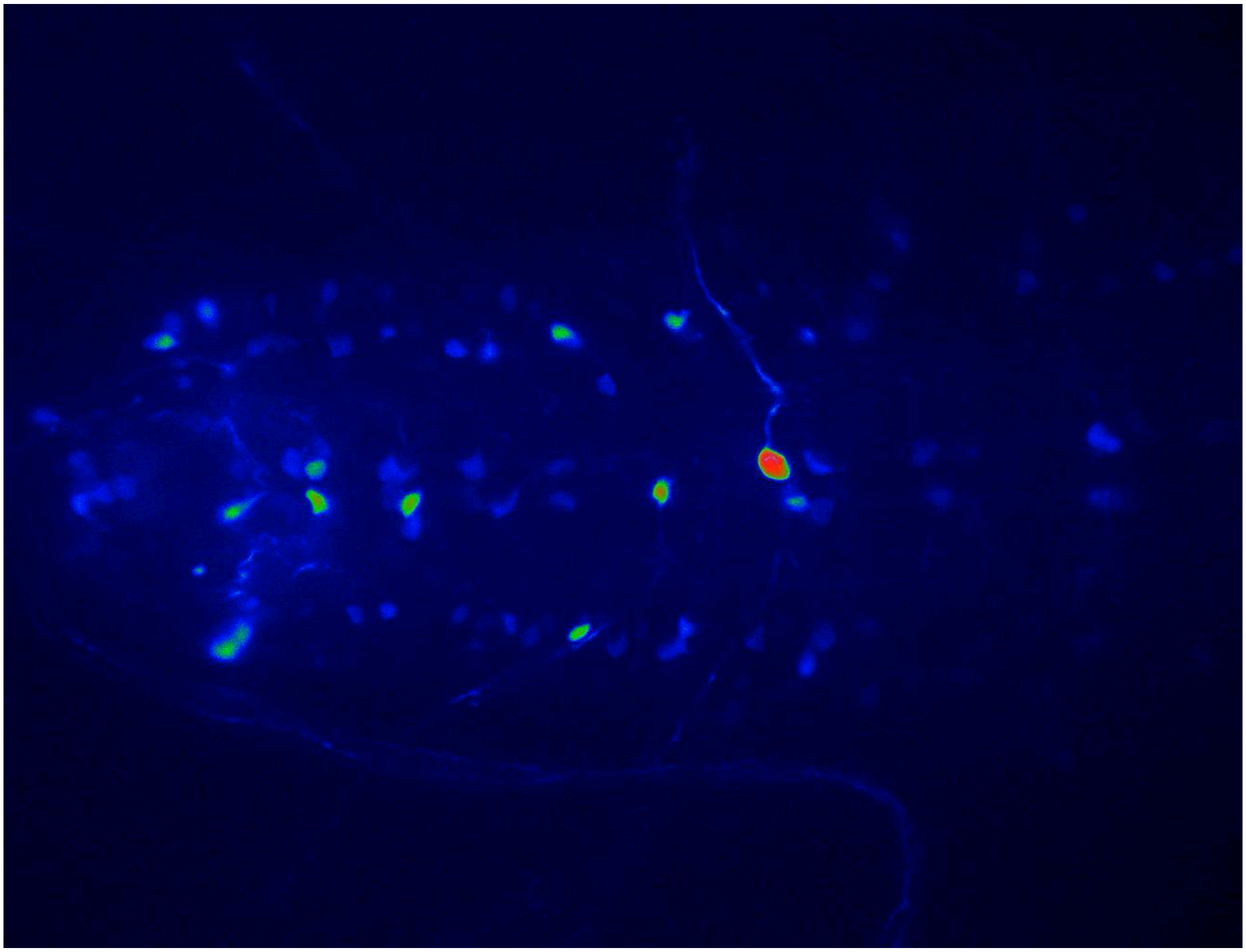

**Supplementary Video 1: Real-time imaging of intracellular calcium signaling in EHR-expressing neurons.**

(A) Example of timecourse of *in vitro* calcium activity in pre-pupal EHR>GCaMP6s CNS) following stimulation with synthetic ETH (600 nM). Images were captured for an hour at a frequency of 0.2 Hz (Video speed 120x). See Fig. 7 for details.

**Table S1: List of all the strains used and their source.**

| GENOTYPE | DESCRIPTION | SOURCE[1] |
|---|---|---|
| $w^{1118}$ (control) | *white* mutant; genetic background of most transgenic animals used here. | Lab stock |
| UAS-EHR (CG10738) | EH Receptor | This study |
| EHR-GAL4 (CG10738) | GAL4 driver for EH Receptor | This study |
| *Tubulin*-GAL4 | GAL4 driver for the microtubule tubulin | BL5138 |
| *n-syb*-GAL4 | GAL4 driver for all neurons neuronal synaptobrevin | BL51942 |
| *eth*-GAL4 | GAL4 driver for ETH (Inka cells) | Ewer lab |
| *386Y*-GAL4 | GAL4 driver for peptidergic and secretory cells | Supplied by P. Taghert |
| *burs*-GAL4 | GAL4 driver for bursicon peptide | Lab stock |
| $EH^{ups}$-GAL4 | GAL4 driver for EH peptide (Vm neurons) | Supplied by S. McNabb |
| $EH^{pan}$-GAL4 | GAL4 driver for Vm and new EH neurons | Supplied by B. White |
| *Mimic{ETHR}*-GAL4 | GAL4 driver for ETH Receptor | BL32718 |
| *vGlut*-GAL4 | GAL4 driver for Vesicular glutamate transporter 1 | BL60312 |
| *tsh*-GAL4 | GAL4 driver for t-shirt (trunk patterning expression) | Supplied by C. Wegener |
| *btl*-GAL4 | GAL4 driver for breathless (tracheal cells) | BL8807 |
| *ChAT*-GAL4 | GAL4 driver for cholinergic neurons | BL60317 |
| *c929*-GAL4 | GAL4 driver for peptidergic neurons | Lab stock |
| *repo*-GAL4 | GAL4 driver for glial cells | Supplied by R. Jackson |
| *c164*-GAL4 | GAL4 driver for motoneurons and other neurons | Supplied by V. Budnik |
| *DDC*-GAL4 | GAL4 driver for dopaminergic and serotonergic neurons | BL7009 |
| *CCAP*-GAL4 | GAL4 driver for CCAP peptide | Ewer lab |
| *trhn*-GAL4 | GAL4 driver for serotoninergic cells | BL84694 |
| *ple*-GAL4 | GAL4 driver for dopaminergic cells | BL86289 |
| *ppk(III)*-GAL4 | GAL4 driver for class IV nociceptive cells | BL32079 |
| *sNPF*-GAL4 | GAL4 driver for sNPF peptide | BL84706 |
| *CCAP-R*-GAL4 | GAL4 driver for CCAP-Receptor | Supplied by B. White |
| *rk*-GAL4 | GAL4 driver for bursicon receptor (*ricket*) | Supplied by B. White |
| *GAD1*-GAL4 | GAL4 driver for glutamatergic cells | Supplied by P. Taghert |
| *nompC*-GAL4 | GAL4 driver for peripheral sensory neurons | Supplied by B. White |
| *Gr66a*-GAL4 | GAL4 driver for gustatory neurons | BL28801 |
| *109(2)80*-GAL4 | GAL4 driver for dendritic neurons, oenocytes and chordotonal organs | BL8769 |
| *Ir20a*-GAL4 | GAL4 driver for gustatory neurons and adult abdominal segments | BL60694 |
| *Ir8a*-GAL4 | GAL4 driver for ionotropic glutamate receptor | BL41731 |
| *Smid c161*-GAL4 | GAL4 driver for Bolwig's nerve, chordotonal organs and imaginal discs) | BL27893 |
| *Ir25a*-GAL4 | GAL4 driver for sensory neurons | BL41728 |
| *Ir7g*-GAL4 | GAL4 driver for gustatory receptor neurons | BL81223 |
| *Ir40a*-GAL4 | GAL4 driver for hydrosensory neurons | BL41727 |

| | | |
|---|---|---|
| EHR-Gal4DBD or -p65AD | Split-GAL4 driver for EHR | This study |
| $EH^{pan}$-p65AD | Split-GAL4 driver for EH peptide (Vm and new targets) | Supplied by B. White |
| ETHR-p65AD | Split-GAL4 driver for ETH Receptor | Supplied by B. White |
| ETHRA-p65AD | Split-GAL4 driver for ETHR isoform A | Supplied by B. White |
| ETHRB-p65AD | Split-GAL4 driver for ETHR isoform B | Supplied by B. White |
| Tub-DBD or -AD | Split-GAL4 driver for microtubule tubulin | BL60298 |
| Tub-DBD or -AD | Split-GAL4 driver for microtubule tubulin | BL60295 |
| *elav*-AD | Split-GAL4 driver for all neurons | Supplied by B. White |
| CCAP-Gal4DBD | Split-GAL4 driver for CCAP peptide | Supplied by B. White |
| CCAP-R-p65AD | Split-GAL4 driver for CCAP receptor | Supplied by B. White |
| *burs*-DBD | Split-GAL4 driver for bursicon peptide | Supplied by B. White |
| *rk*-AD | Split-GAL4 driver for bursicon receptor | Supplied by B. White |
| vGlut-p65AD | Split-GAL4 driver for vesicular glutamate transporter 1 | Supplied by B. White |
| ChAT-p65AD | Split-GAL4 driver for choline acetyltransferase | Supplied by B. White |
| Tub-GAL80ts | GAL80 line temperature sensitive | Lab stock |
| UAS-EHR RNAi | RNAi against EHR | BL38346 |
| UAS-EHR RNAi | RNAi against EHR | BL57318 |
| UAS-EHR RNAi | RNAi against EHR | BL60439 |
| UAS-EHR RNAi | RNAi against EHR | BL28580 |
| UAS-EHR RNAi | RNAi against EHR | NIG10738-R1, III |
| UAS-EHR RNAi | RNAi against EHR | NIG10738-R2, II |
| UAS-ETHR RNAi | RNAi against ETHR | VDRC42717 |
| *Df(3L)exel9017* | Genetic deletion that includes CG10738 | BL7934 |
| Minos(EHR) | Minos insertion in CG10738 gene | BL24564 |
| Mimic(EHR) | Mimic insertion in CG10738 gene | BL35974 |
| UAS-GFP | Green Fluorescent Protein (GFP) | BL6874 |
| UAS-GFP nuclear | Nuclear GFP | BL4775 |
| UAS-RFP nuclear | Nuclear Red Fluorescent Protein (RFP) | Lab stock |
| UAS-mCherry | Monomeric cherry fluorescent protein | BL27391 |
| UAS-esg-GFP | escargot (esg)-GFP fusion protein | Supplied by B. White |
| UAS-*reaper* | Apoptotic factor | BL5824 |
| UAS-*Kir.2.1* | Inwardly rectifying potassium channel | Supplied by B. White |
| UAS-GCaMP6s | Calcium sensitive GFP | BL42749 |
| UAS-*TrpA1* | Transient receptor potential cation channel | BL26263 |
| UAS-d*cr2* | Dicer-2 | Lab stock |

[1]BL: Bloomington *Drosophila* stock center (Bloomington, USA); NIG: Fly stocks of National Institute of Genetics (Mishima, Japan); VDRC: Vienna Drosophila Resource Center (Vienna, Austria).

**Table S2: EHR RNAi screening**. Phenotype of RNAi lines for EHR expressed using *tubulin*-GAL4.

| RNAi line | Phenotype |
|---|---|
| BL38346 (Bloomington, USA) | Normal pupal ecdysis. Some adults with wing expansion defects |
| BL57318 (Bloomington, USA) | Crosses produced few animals; no adult eclosed |
| BL60439 (Bloomington, USA) | Some pupae with ecdysial problems |
| BL28580 (Bloomington, USA) | Pupae with ecdysial problems; no adult eclosed |
| CG10738 RNAI-R1(NIG, Japan) [1] | Pupae with ecdysial problems; no adult eclosed |
| CG10738 RNAI-R2 (NIG, Japan) | Normal ecdyses |

[1] Strongest line; used for results reported here.

### Table S3: Summary of statistical analyses

| Fig. | Test | Conditions Analyzed | P value | P < 0.05) |
|---|---|---|---|---|
| 1F | Chi-square test | $w^{1118}$ control *vs* $w^{1118}$ +EH | <0,0001 | Yes |
| 1F | Chi-square test | $w^{1118}$ control *vs Mi{EHR}/Df(3)EHR* + EH | 0,1292 | No |
| 1F | Chi-square test | $w^{1118}$ control *vs EHR*-GAL4/*Df(3)EHR* + EH | 0,0021 | Yes |
| 1F | Chi-square test | $w^{1118}$ +EH *vs Mi{EHR}/Df(3)EHR* + EH | <0,0001 | Yes |
| 1F | Chi-square test | $w^{1118}$ +EH vs *EHR*-GAL4/*Df(3)EHR* + EH | <0,0001 | Yes |
| 1F | Chi-square test | *Mi{EHR}/Df(3)EHR* + EH vs *EHR*-GAL4/*Df(3)EHR* + EH | 0,1277 | No |
| 1G | Chi-square test | $w^{1118}$ dVP+30 vs *Mi{EHR}/Df(3)EHR* dVP +2h | <0,0001 | Yes |
| 1G | Chi-square test | $w^{1118}$ dVP+30 vs *EHR*-GAL4/*Df(3)EHR* dVP +2h | <0,0001 | Yes |
| 1G | Chi-square test | *Mi{EHR}/Df(3)EHR* dVP +2h vs *EHR*-GAL4/*Df(3)EHR* dVP +2h | <0,0001 | Yes |
| 2A | Unpaired t test | $w^{1118}$ *vs Df(3)EHR/Mi{EHR}* - Locomotion pre-TC | <0,0001 | Yes |
| 2A | Unpaired t test | $w^{1118}$ *vs Df(3)EHR/Mi{EHR}* - Locomotion after-TC | 0,0012 | Yes |
| 2A | Unpaired t test | $w^{1118}$ *vs Df(3)EHR/Mi{EHR}* - pre-ecdysis | N/A | N/A |
| 2A | Unpaired t test | $w^{1118}$ *vs Df(3)EHR/Mi{EHR}* - ecdysis | 0,0289 | Yes |
| 2A | Unpaired t test | $w^{1118}$ *vs Df(3)EHR/EHR*-GAL4 - Locomotion pre-TC | <0,0001 | Yes |
| 2A | Unpaired t test | $w^{1118}$ *vs Df(3)EHR/EHR*-GAL4 - Locomotion after TC | 0,0187 | Yes |
| 2A | Unpaired t test | $w^{1118}$ *vs Df(3)EHR/EHR*-GAL4 - pre-ecdysis | N/A | N/A |
| 2A | Unpaired t test | $w^{1118}$ *vs Df(3)EHR/EHR*-GAL4 - ecdysis | <0,0001 | Yes |
| 2A | Unpaired t test | $w^{1118}$ *vs UAS-EHR;EHR*-GAL4>*Df(3)EHR* - Locomotion pre-TC | 0,6559 | No |
| 2A | Unpaired t test | $w^{1118}$ *vs UAS-EHR;EHR*-GAL4>*Df(3)EHR* - Locomotion after TC | 0,949 | No |
| 2A | Unpaired t test | $w^{1118}$ *vs UAS-EHR;EHR*-GAL4>*Df(3)EHR* - pre-ecdysis | 0,0016 | Yes |
| 2A | Unpaired t test | $w^{1118}$ *vs UAS-EHR;EHR*-GAL4>*Df(3)EHR* - ecdysis | 0,0445 | No |
| 2A | Unpaired t test | *Df(3)EHR/Mi{EHR} vs Df(3)EHR/EHR*-GAL4 - Locomotion pre-TC | 0,153 | No |
| 2A | Unpaired t test | *Df(3)EHR/Mi{EHR} vs Df(3)EHR/EHR*-GAL4 - Locomotion after TC | 0,2423 | No |
| 2A | Unpaired t test | *Df(3)EHR/Mi{EHR} vs Df(3)EHR/EHR*-GAL4 - pre-ecdysis | N/A | N/A |
| 2A | Unpaired t test | *Df(3)EHR/Mi{EHR} vs Df(3)EHR/EHR*-GAL4 - ecdysis | 0,0666 | No |
| 2A | Unpaired t test | *Df(3)EHR/Mi{EHR} vs UAS-EHR;EHR*-GAL4>*Df(3)EHR* - Locomotion pre-TC | <0,0001 | Yes |
| 2A | Unpaired t test | *Df(3)EHR/Mi{EHR} vs UAS-EHR;EHR*-GAL4>*Df(3)EHR* - Locomotion after TC | <0,0001 | Yes |
| 2A | Unpaired t test | *Df(3)EHR/Mi{EHR} vs UAS-EHR;EHR*-GAL4>*Df(3)EHR* - pre-ecdysis | N/A | N/A |
| 2A | Unpaired t test | *Df(3)EHR/Mi{EHR} vs UAS-EHR;EHR*-GAL4>*Df(3)EHR* - ecdysis | 0,0005 | Yes |

| | | | | |
|---|---|---|---|---|
| 2A | Unpaired t test | *Df(3)EHR/EHR*-GAL4 *vs* *UAS-EHR;EHR*-GAL4>*Df(3)EHR* - Locomotion pre-TC | <0,0001 | Yes |
| 2A | Unpaired t test | *Df(3)EHR/EHR*-GAL4 *vs* *UAS-EHR;EHR*-GAL4>*Df(3)EHR* - Locomotion after TC | 0,0005 | Yes |
| 2A | Unpaired t test | *Df(3)EHR/EHR*-GAL4 *vs* *UAS-EHR;EHR*-GAL4>*Df(3)EHR* - pre-ecdysis | N/A | N/A |
| 2A | Unpaired t test | *Df(3)EHR/EHR*-GAL4 *vs* *UAS-EHR;EHR*-GAL4>*Df(3)EHR* - ecdysis | <0,0001 | Yes |
| 2D | One-way ANOVA + Tukey's | $w^{1118}$ vs. *Mi{EHR}/Df(3)EHR* - Duration of tracheal air filling | <0,0001 | Yes |
| 2D | One-way ANOVA + Tukey's | $w^{1118}$ vs. *EHR*-GAL4*/Df(3)EHR* - Duration of tracheal air filling | <0,0001 | Yes |
| 2D | One-way ANOVA + Tukey's | $w^{1118}$ vs. *UAS-EHR; EHR*-GAL4*/Df(3)EHR* - Duration of tracheal air filling | 0,9972 | No |
| 2D | One-way ANOVA + Tukey's | *Mi{EHR}/Df(3)EHR* vs. *EHR*-GAL4*/Df(3)EHR* - Duration of tracheal air filling | 0,9975 | No |
| 2D | One-way ANOVA + Tukey's | *Mi{EHR}/Df(3)EHR* vs. *UAS-EHR; EHR*-GAL4*/Df(3)EHR* - Duration of tracheal air filling | <0,0001 | Yes |
| 2D | One-way ANOVA + Tukey's | *EHR*-GAL4*/Df(3)EHR* vs. *UAS-EHR; EHR*-GAL4*/Df(3)EHR* - Duration of tracheal air filling | <0,0001 | Yes |
| 3A | Unpaired t test | $w^{1118}$>*EHR*-GAL4 vs $w^{1118}$>2x*Kir2.1* - Locomotion pre-TC | 0,7721 | No |
| 3A | Unpaired t test | $w^{1118}$>*EHR*-GAL4 vs $w^{1118}$>2x*Kir2.1* - Locomotion after TC | 0,8184 | No |
| 3A | Unpaired t test | $w^{1118}$>*EHR*-GAL4 vs $w^{1118}$>2x*Kir2.1* - pre-ecdysis | 0,9387 | No |
| 3A | Unpaired t test | $w^{1118}$>*EHR*-GAL4 vs $w^{1118}$>2x*Kir2.1* - ecdysis | 0,7045 | No |
| 3A | Unpaired t test | $w^{1118}$> *EHR*-GAL4 vs *EHR*-GAL4>2x*Kir2.1* - Locomotion pre-TC | <0,0001 | Yes |
| 3A | Unpaired t test | $w^{1118}$>*EHR*-GAL4 vs *EHR*-GAL4>2x*Kir2.1* - Locomotion after TC | <0,0001 | Yes |
| 3A | Unpaired t test | $w^{1118}$>*EHR*-GAL4 vs *EHR*-GAL4>2x*Kir2.1* - pre-ecdysis | 0,0091 | Yes |
| 3A | Unpaired t test | $w^{1118}$>*EHR*-GAL4 vs *EHR*-GAL4>2x*Kir2.1* - ecdysis | <0,0001 | Yes |
| 3A | Unpaired t test | $w^{1118}$>2x*Kir2.1* vs *EHR*-GAL4>2x*Kir2.1* - Locomotion pre-TC | 0,0005 | Yes |
| 3A | Unpaired t test | $w^{1118}$>2x*Kir2.1* vs *EHR*-GAL4>2x*Kir2.1* - Locomotion after TC | <0,0001 | Yes |
| 3A | Unpaired t test | $w^{1118}$>2x*Kir2.1* vs *EHR*-GAL4>2x*Kir2.1* - pre-ecdysis | 0,014 | Yes |
| 3A | Unpaired t test | $w^{1118}$>2x*Kir2.1* vs *EHR*-GAL4>2x*Kir2.1* - ecdysis | <0,0001 | Yes |
| 3A | Unpaired t test | $w^{1118}$>*rpr* vs *EHR*-GAL4>*rpr* - Locomotion pre-TC | 0,1235 | No |
| 3A | Unpaired t test | $w^{1118}$>*rpr* vs *EHR*-GAL4>*rpr* - Locomotion after TC | <0,0001 | Yes |
| 3A | Unpaired t test | $w^{1118}$>*rpr* vs *EHR*-GAL4>*rpr* - pre-ecdysis | N/A | N/A |
| 3A | Unpaired t test | $w^{1118}$>rpr vs *EHR*-GAL4>*rpr* - ecdysis | <0,0001 | Yes |
| 7E | LME model | Rest population mean time to peak | 0.052509 | No |
| 7E | LME model | CCAP Left population mean time to peak | 0.037704 | Yes |
| 7E | LME model | CCAP Right population mean time to peak | 0.1723 | No |

| | | | | |
|---|---|---|---|---|
| 7E | LME model | Central Right population mean time to peak | 0.83524 | No |
| 7E | LME model | Central Left population mean time to peak | 0.84542 | No |
| 7E | LME model | Posterior Right population mean time to peak | 0.71989 | No |
| 7F | Bonferroni + Wilcoxon rank | Rest population vs CCAP Left population | 0.438837 | No |
| 7F | Bonferroni + Wilcoxon rank | Rest population vs CCAP Right population | 0.378358 | No |
| 7F | Bonferroni + Wilcoxon rank | Rest population vs Central Left population | 1.000000 | No |
| 7F | Bonferroni + Wilcoxon rank | Rest population vs Central Right population | 0.671896 | No |
| 7F | Bonferroni + Wilcoxon rank | Rest population vs Posterior (POS) population | 1.000000 | No |
| 7F | Bonferroni + Wilcoxon rank | CCAP Left population vs CCAP Right population | 1.000000 | No |
| 7F | Bonferroni + Wilcoxon rank | CCAP Left population vs Central Right population | 0.007175 | Yes |
| 7F | Bonferroni + Wilcoxon rank | CCAP Left population vs Central Left population | 0.000217 | Yes |
| 7F | Bonferroni + Wilcoxon rank | CCAP Left population vs Posterior (POS) population | 0.021207 | Yes |
| 7F | Bonferroni + Wilcoxon rank | CCAP Right population vs Central Left population | 0.025342 | Yes |
| 7F | Bonferroni + Wilcoxon rank | CCAP Right population vs Central Right population | 0.000517 | Yes |
| 7F | Bonferroni + Wilcoxon rank | CCAP Right population vs Posterior (POS) population | 0.020122 | Yes |
| 7F | Bonferroni + Wilcoxon rank | Central Left population vs Central Right population | 1.000000 | No |
| 7F | Bonferroni + Wilcoxon rank | Central Left population vs Posterior (POS) population | 1.000000 | No |
| 7F | Bonferroni + Wilcoxon rank | Central Right population vs Posterior (POS) population | 1.000000 | No |
